# A distinct set of brain areas process prosody—the melody of speech

**DOI:** 10.64898/2025.12.12.693781

**Authors:** Tamar I. Regev, Hee So Kim, Niharika Jhingan, Sara Swords, Hope Kean, Colton Casto, Jennifer S. Cole, Evelina Fedorenko

## Abstract

Human speech carries information beyond the words themselves: pitch, loudness, duration, and pauses—jointly referred to as ‘prosody’—emphasize critical words, help group words into phrases, and convey emotional and other socially-relevant information. Using a novel fMRI paradigm, we identify a set of prosody-responsive brain areas and then richly characterize them across 8 experiments (25 experimental conditions; n=51 participants). These areas—located on the lateral temporal surface bilaterally and in the frontal lobe—respond to the presence of prosody in both meaningful and meaningless speech, and are distinct from nearby temporal pitch- and speech-perception areas and from frontal areas sensitive to general cognitive and attentional demands. They are also dissociable from—but show partial overlap with—the language areas and with the area that processes dynamic facial expressions. Thus, prosodic information is processed by a distinct set of brain areas, which may help integrate linguistic meanings with non-verbal communicative signals.

## 1. INTRODUCTION

When we speak, we transmit information not only using words, but also via non-verbal communication channels including facial expressions, gestures, and speech prosody (Goldin-Meadow, 2014; Vigliocco et al., 2014; Holler & Levinson, 2019; Hadley et al., 2022; Hagoort & Özyürek, 2024). Here, we focus on prosody, which refers to the melodic and rhythmic patterning in speech at the phrase level—encoding information beyond the words themselves. Time-varying patterns of pitch and loudness in speech, along with local tempo variation (faster or slower articulation rate) and pauses convey a wealth of information about the structure and meaning of an utterance (Wagner & Watson, 2010; Cole, 2015; Tomasello et al., 2022; Ladd & Arvaniti, 2023). For example, prosody can transform a declarative sentence (a statement) into a question (Hellbernd & Sammler, 2016), indicate how words are grouped into phrases (Degano et al. 2024; Breen et al. 2011; Snedeker & Trueswell, 2003; Watson & Gibson, 2004), mark words that are most important in an utterance (Breen et al., 2010; Roettger et al., 2019), and express speaker attitudes such as sarcasm, or humorous tone (Cheang & Pell, 2008; Menninghaus et al., 2014). Prosodic cues also convey speaker emotions (Ladd et al., 1985; Pell et al., 2009; Larrouy-Maestri et al., 2025) and information relevant to the social dynamics (Heldner et al., 2010; Van Zyl et al., 2013). However, although prosody is an integral component of human communication, its neural basis remains debated (Wildgruber et al., 2006; Belyk & Brown, 2014; Themistocleous, 2025).

*A priori*, where in the brain might we look for responses to prosody perception? Prosody falls at the intersection of auditory perception, language processing, and social perception and reasoning. Past research has revealed that many components of these abilities rely on specialized neural machinery, including brain areas selective for processing pitch (Norman-Haignere et al., 2013), for processing speech sounds (Norman-Haignere et al., 2015; Overath, McDermott, et al., 2015), for language comprehension (Fedorenko et al., 2024), for different aspects of visual and auditory social perception (Kanwisher et al., 1997; Belin, Zatorre, Lafalli, et al., 2000; Downing et al., 2001; Pitcher et al., 2011; Deen et al., 2015; Isik et al., 2017; Papeo & Abassi, 2019), and for social reasoning (Fletcher et al., 1995; Gallagher et al., 2000; Vogeley et al., 2001; Ruby & Decety, 2003; Saxe & Kanwisher, 2003; Kandylaki et al., 2015). Does interpreting prosodic patterns—which are rich in pitch changes, intimately tied to the phonological content of the speech signal, and often critical to understanding linguistic messages and reasoning about speaker intent—draw on any of these established mechanisms? Or does prosody perception instead rely on its own specialized neural system?

Past work has left this question open. Most prior studies have relied on methods with relatively poor spatial resolution such as investigations of patients with brain damage (Ross, 1981; Lancker & Sidtis, 1992; Pell & Baum, 1997; Walker et al., 2002; Kucharska-Pietura et al., 2003; for reviews see Witteman et al., 2011; Hu et al., 2025), ERPs (Steinhauer et al. 1999; 2001; Steinhauer 2003; Paulmann & Kotz 2008; Paulmann et al. 2012), and group-averaged fMRI (Meyer et al., 2002; Plante et al., 2002, 2006; Sammler et al., 2015; Seydell-Greenwald et al., 2020; for reviews see Wildgruber et al., 2006; Belyk & Brown, 2014), which blurs functional responses due to well-established inter-individual variability in the precise locations of functional areas (Frost & Goebel, 2012; Tahmasebi et al., 2012; Fedorenko, 2021; Du et al., 2024). Many of these studies have implicated lateral temporal and frontal cortical areas (Belyk & Brown, 2014), often arguing for a right-hemisphere bias for emotional prosody perception (Alba-Ferrara et al., 2012; Buchanan et al., 2000; Mitchell et al., 2003; Seydell-Greenwald et al., 2020; Wildgruber et al., 2006; cf. Kotz et al. 2003). Some have also reported a right-hemisphere bias for linguistic prosody perception (Sammler et al., 2015), but others have observed left-dominant or bilateral activations (van der Burght et al., 2019;Kreitewolf et al., 2014). Some studies have reported cerebellar and subcortical contributions to prosody processing (Pell & Leonard, 2003; Friederici et al., 2007; Paulmann et al., 2008; Sammler et al., 2010; Adamaszek et al., 2014). A challenge in interpreting these past studies with respect to the nature of prosodic representations and computations is that 1) they have typically used a single paradigm each, and 2) the broad reported anatomical areas are highly functionally heterogeneous. For example, many of the specialized areas noted above tile the lateral surface of the temporal lobes. Do the previously reported temporal prosody-responsive areas overlap with the pitch-processing areas? With the speech perception areas? With the language areas? With areas that process dynamic faces? Without comparing between tasks that engage those different functions within individuals, it is impossible to determine.

To systematically characterize prosody-responsive brain areas, we turned to the functional localization approach, which has been successful in other domains (Kanwisher, 2010). Using a novel paradigm for detecting sensitivity to prosody, we identified a set of temporal and frontal brain areas, and then examined their responses to diverse conditions and their overlap with known functional areas. We found that prosody is processed by brain areas distinct from low-level auditory areas and from areas sensitive to general attentional demands but partially overlapping with the language network—a set of brain areas that support word recognition and word combinatorial linguistic processing (Fedorenko et al., 2024)—and areas that process dynamic facial expressions (Pitcher et al., 2011; Pitcher & Ungerleider, 2021; Prabhakar et al., 2025). The overlap between prosody areas and other communication-relevant mechanisms may arise from the need to integrate information from different communication channels during face-to-face interaction, and may suggest even greater overlap in pre-linguistic infants, who rely exclusively on non-verbal cues.

## 2. RESULTS

To identify brain areas that are sensitive to prosody, we contrasted sentences uttered with expressive, natural prosody (Sentences Prosody+, or SP+) with the same sentences presented with disrupted prosody that lacks linguistic or social-emotional cues typically present in speech (Sentences Prosody-, or SP-). For SP+, we extracted sentences from conversational corpora of American English (Switchboard and COCA; Davies 2010), and had an actress record them using natural expressive prosody. To generate SP-, the same actress recorded the individual words in isolation, which were then combined into sentences with disrupted prosodic contours (**Figure 1A**, <u>Methods</u>). We reasoned that prosody-responsive brain areas (henceforth *prosody areas*) should respond more strongly to the sentences with expressive prosody relative to the sentences with disrupted prosody (SP+ > SP-), reflecting sensitivity to natural prosodic patterns.

Furthermore, prosody can convey information even when the meaning of the words is inaccessible. For example, infants rely on prosody during language learning (Piazza et al. 2017; Saint-Georges et al. 2013; Fernald 1985; Furrow et al. 1979), and adults can infer affective states and communicative intent from speech in an unfamiliar language or muffled conversations heard from far away (Pell & Skorup, 2008; Paulmann & Uskul, 2014). We therefore created nonword analogs of the SP+ and SP-conditions (<u>Methods</u>, **Figure 1A**) and reasoned that prosody brain areas should respond more strongly to the nonword sentences with expressive prosody relative to those with disrupted prosody (NP+ > NP-). Thus, the critical contrast for identifying prosody brain areas (termed the *prosody localizer contrast*) was a conjunction of the SP+>SP- and the NP+>NP-contrasts (<u>Methods</u>; all stimuli are available at OSF: https://osf.io/2t9j3/). This contrast was designed to capture both linguistic and emotional aspects of prosody.

Finally, as an acoustic control, we temporally inverted the NP+ and NP– stimuli to create two additional conditions (InvP+ and InvP–, **Figure 1A**). The inversion manipulation destroys natural prosody, while preserving low-level sound properties, so we expected minimal responses to these conditions in prosody areas. Because we were interested in the brain mechanisms that underlie naturalistic (rather than goal-/task-driven) prosodic processing, participants were simply instructed to listen attentively and to press a button when they see a picture of a finger pressing a button (<u>Methods</u>).

Fifty-one unique participants (mean age = 24.8 years, st. dev. = 7 years; 29 females; **Table 1**, <u>Methods</u>) were recruited. All participants performed the prosody localizer task, and each participant performed some additional tasks with the goal of characterizing the functional profiles of the prosody areas and comparing them to known functional systems.

### 2.1 A set of bilateral brain areas—most prominently in the temporal lobes—respond to prosody

The prosody contrast elicited a response in a set of temporal and frontal areas in the left and the right hemispheres (**Figure 1B**). We used the group-constrained subject-specific (GSS, <u>Methods</u>) functional localization approach (Fedorenko et al., 2010; Julian et al., 2012), which searches for areas that are spatially consistent across participants, but allows for inter-individual variability in the precise locations of functional areas. In particular, a group-level probabilistic overlap map for the relevant contrast (**Figure 1E**) is divided into ‘parcels’—areas of most spatially consistent activation. Parcels that are sufficiently large and show activity in most participants (more than 90%, **Figure S1**) are selected for further analyses, where they are used to constrain the definition of individual-level functional regions of interest (fROIs). fROIs are defined as the top 10% of most localizer-responsive voxels within each parcel (**Figure 1D**) using a portion of the data, to maintain independence from the data used for response estimation (Kriegeskorte et al., 2009).

The parcels that emerged from the GSS analysis can be grouped into three sets: lateral temporal areas (‘temporal’), lateral frontal areas (‘lateral-frontal’), and medial frontal areas (‘medial-frontal’). All three sets showed (in left-out data) robust sensitivity to prosody in both meaningful stimuli (SP+>SP-; *p*s<0.05, linear mixed-effects (LME) model, **Figure 1B**, **Table S1**) as well as meaningless, nonword stimuli (NP+>NP-; *p*s<0.05, **Figure 1B**, **Table S1**; see **Figure S2** for the functional response profiles of the individual areas). The responses to the control conditions (InvP+ and InvP-) were low, and not different from each other across the fROIs (**Table S1**), ruling out the possibility of low-level acoustic differences driving the prosody responses. Interestingly, prosody areas showed a main effect of intelligibility (SP+>NP+ and SP->NP-; **Table S2**), which suggests that these areas are sensitive to verbal content in addition to prosody; we return to this point in Section 2.4.

To ensure that the responses in the prosody areas generalize beyond the particular stimuli and speaker used in the localizer experiment, a subset of participants (n=24; Experiment 2, <u>Methods</u>) listened to a more diverse set of recordings of natural speech (in American English) from multiple speakers, extracted from a corpus of phone-call conversations (CALLFRIEND, 2^nd^ edition, Canavan et al. 2019). These sentences were similar to those in the SP+ condition except that we used the original productions rather than their reproductions by an actress. We additionally included a different set of nonword sentences produced with natural spontaneous prosody (<u>Methods</u>), similar to the NP+ condition in the localizer experiment, and a condition where the phone-call stimuli were played backwards, similar to the InvP+ control condition. In the temporal prosody areas, the responses to these three conditions were similar to the corresponding conditions in the prosody localizer, establishing the robustness of the responses to variation in the experimental materials (**Figure 2B**, Experiment 2 panel). The frontal prosody areas responded only weakly to these conditions, indicating poorer generalizability across different kinds of prosody-rich stimuli (**Figure S3**, stimuli available on OSF).

Having identified a set of prosody-responsive areas, we next examined their relationship to known brain areas and networks that i) support perceptual or cognitive processes that bear some relationship to prosodic processing, and ii) have components in the same general anatomical areas where we found responses to prosody. These areas include auditory (pitch and speech) processing areas (Section 2.2), areas of the Multiple Demand network, which are sensitive to general cognitive and attentional demands (Section 2.3), areas of the language network (Section 2.4), and areas that support the processing of dynamic faces (Section 2.5). We primarily focus on the temporal prosody-responsive areas given that they respond most consistently to prosody, but we do consider the frontal areas in Sections 2.3 and 2.4.

**Figure 1.**
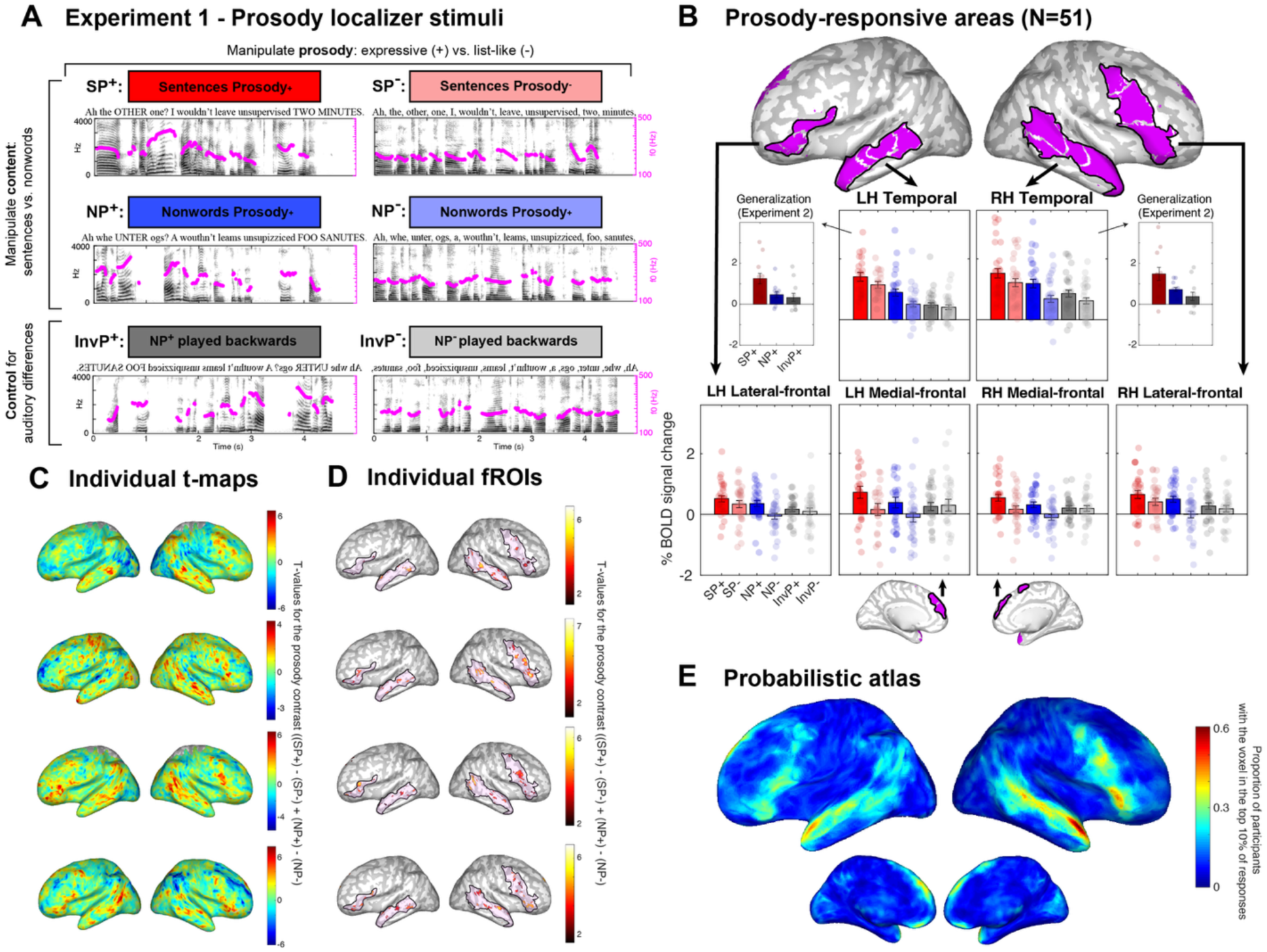
Prosody localizer (Experiment 1) stimuli and results. **A)** Conditions of the fMRI prosody localizer paradigm. Four critical conditions were developed, crossing linguistic content (sentences versus phonologically-matched nonword sentences, where linguistic content is not interpretable) and prosodic expressivity (natural expressive prosody or list-like prosody, atypical of connected speech utterances). The prosody localizer contrast was defined as a conjunction (see <u>Methods</u> for implementation) of the prosody contrasts with or without linguistic content. Two additional conditions were included to control for low-level acoustic differences: the nonword conditions were played backwards, such that neither the content nor the prosody was familiar or meaningful. **B)** The areas that were discovered with the prosody localizer contrast are grouped into three sets (temporal, lateral-frontal and medial-frontal) in each hemisphere, and the mean responses (across all individual areas) of each set to the 6 conditions in A are presented. We show the parcels, which are used to constrain the definition of individual-level fROIs (<u>Methods</u>); the fROIs are 10% of the parcels in size (see panel **D** for examples). For each hemisphere, we show the lateral and medial views (we use surface projection for visualization only; all the analyses were performed in the volume; <u>Methods</u>). The bar graphs represent the average (across participants) responses, in % BOLD signal change relative to the fixation baseline. To each condition extracted from individual-level fROIs (error bars are standard error of the mean across participants, and dots correspond to individual participants). Localization and effect size estimation were performed using independent subsets of the data to avoid circularity (<u>Methods</u>). See **Figure S2** for responses of the individual areas comprising the sets. Insets to the sides of the temporal areas’ profiles show the generalization of these regions’ responses to SP+, NP+, and InvP+ conditions from Experiment 2, which used more diverse stimuli (<u>Methods</u>; see **Figure S3** for responses in the frontal prosody areas, which showed poorer generalization). **C)** Whole brain t-maps for the prosody localizer contrast for 4 sample participants. **D)** Individual-level prosody fROIs for 4 sample participants. These fROIs are defined as the top 10% of prosody-responsive voxels in each parcel (the color scale shows the t-values for the prosody localizer contrast). The black outlines show parcel boundaries (conjunction of contiguous parcels for the temporal and lateral-frontal sets). **E)** Probabilistic activation overlap map for the prosody contrast for all participants (n=51; <u>Methods</u>).

**Figure 2.**
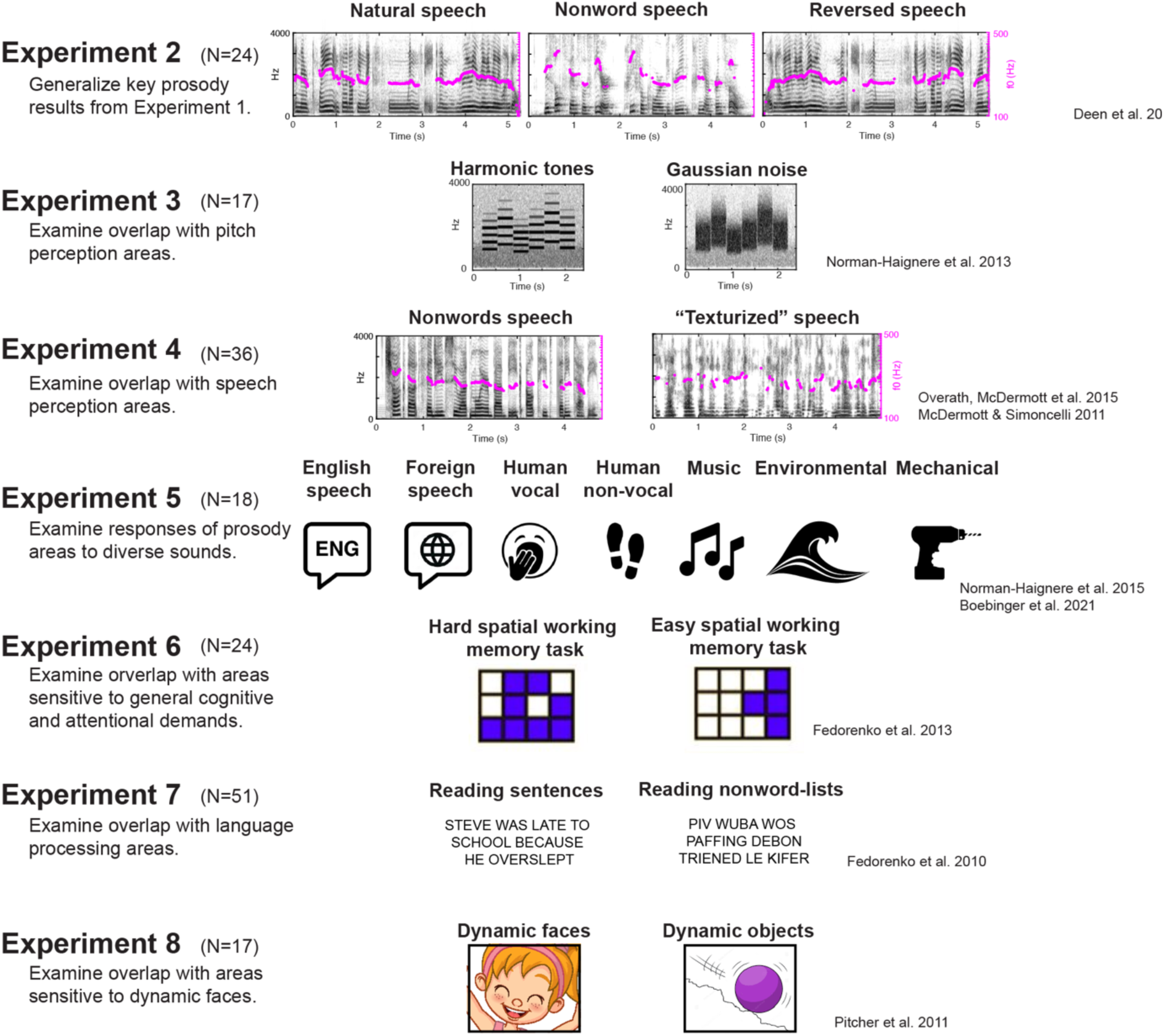
An overview of all experiments beyond the prosody localizer experiment (Experiment 1, Figure 1). For each experiment, we provide the objective, the number of participants (N), the main conditions with schematic illustrations of sample stimuli, and the references to the papers where each paradigm was introduced and/or validated.

### 2.2 Prosody-responsive areas are distinct from auditory (pitch and speech perception) areas

We first asked whether the prosody areas in the temporal lobe are distinct from areas that specialize for auditory functions that are relevant to prosody: namely, pitch perception and speech perception. *Pitch* is a core component of prosody, giving rise to intonation contours that can carry both linguistic and social-emotional information (Hirschberg, 2006; Scherer, 2013; Prieto, 2015; Büring, 2016; Larrouy-Maestri et al., 2025). Furthermore, prosody accompanies—and is inherently tied to—the *speech* signal. Our prosody localizer already controls for the presence of pitch (pitch is present across all six conditions) and for speech sounds (present across both sentence and nonwords conditions). However, here, we more systematically test for overlap between the prosody areas and a) the pitch-processing areas (Norman-Haignere et al., 2013), and b) the areas that support the perception of speech sounds (Overath et al., 2015).

The bilateral pitch-processing area (Norman-Haignere et al., 2013, 2015) responds to harmonic sounds, which elicit the percept of pitch, and is evolutionarily recent (Norman-Haignere et al., 2019). We used an established contrast between harmonic tones and frequency-matched noise to localize this area (Norman-Haignere et al., 2013; **Figure 2**, Experiment 3). The bilateral speech-processing area responds selectively to the acoustic features of human speech regardless of whether the content is intelligible (Overath et al., 2015; Norman-Haignere et al., 2015). We used an established contrast between unintelligible speech recordings (nonwords here) and acoustically-matched synthesized control sounds (‘Speech textures’; McDermott & Simoncelli, 2011) that are not perceived as speech (Overath et al., 2015; **Figure 2**, Experiment 4; <u>Methods</u>).

Replicating prior studies (Norman-Haignere et al., 2013; Overath et al., 2015), the pitch and speech contrasts elicited robust responses in bilateral temporal areas within primary and secondary auditory cortex (**Figure 3B-C**). Critically, both the pitch and speech areas responded to the conditions of the prosody experiment in a distinct way from the prosody areas (**Figure 3A-C**). The pitch areas responded strongly and uniformly across conditions (**Figure 3B**), consistent with the fact that all stimuli used harmonic sounds and evoked a pitch percept. The lack of an overall reduced response to the Prosody-conditions compared to the Prosody+ conditions (p>0.05, LME, <u>Methods</u>) suggests that the pitch area is not strongly sensitive to the pitch modulations that unfold over time in prosodic contours, plausibly because this area’s temporal integration window is short (∼0.5s; Norman-Haignere et al., 2022) relative to prosodic modulations at the phrase level which plausibly have a longer time scale. The speech areas’ responses were also not modulated by prosody (**Figure 3C**, p>0.05, <u>Methods</u>). Further, in contrast to the prosody areas, the speech areas’ responses were only weakly modulated by intelligibility, consistent with previous findings (Norman-Haignere et al., 2015) (p=0.043, LME comparing all sentences to all nonwords in the prosody task, <u>Methods</u>; note that this intelligibility effect was driven by the Prosody-conditions, **Figure 3C**). Finally, the two control conditions (InvP+ and InvP-) elicited lower responses than the prosody conditions in the speech areas (p<0.001, LME comparing the Prosody+ and Prosody-conditions to the control conditions: InvP+ and InvP-), plausibly because some acoustic characteristics of reversed speech make it sound less ‘speech-like’.

The prosody areas responded similarly weakly to both conditions of the pitch localizer (Experiment 3), indicating lack of sensitivity to the mere presence of pitch (p>0.05, **Table S3**). For the speech conditions (Experiment 4), the prosody-responsive areas did show a higher response to nonword speech (which was characterized by natural-like prosody) compared to speech textures (p<0.01, **Table S3**), although this effect was substantially weaker than in the speech areas (p<0.001, interaction term of the speech localizer effect between the prosody and the speech areas, **Table S5**, <u>Methods</u>). The reliable response in the prosody areas to nonword speech provides yet another generalization to a new set of auditory prosody-rich stimuli.

To complement the univariate analyses, we examined the fine-grained activation patterns across voxels for the prosody, pitch, and speech contrasts within a large temporal lobe mask (**Figure 3D**). For each contrast, we assessed the stability of the activation pattern across runs using a split-half approach, which served as the noise ceiling for the spatial correlations across contrasts (<u>Methods</u>).

The activation pattern was more stable for the speech contrast than for the pitch and prosody contrasts (**Figure 3D**), but critically, the activation patterns between the prosody and non-prosody (pitch or speech) contrasts were less similar compared to the similarity of voxel activation patterns within tasks across runs (p<0.001 both for pitch and speech versus prosody, LME, Methods, **Figure 3D, Table S6**).

The univariate and multivariate analyses therefore converge to suggest that the prosody areas are spatially and functionally distinct from the auditory areas that specialize for pitch and speech perception.

**Figure 3.**
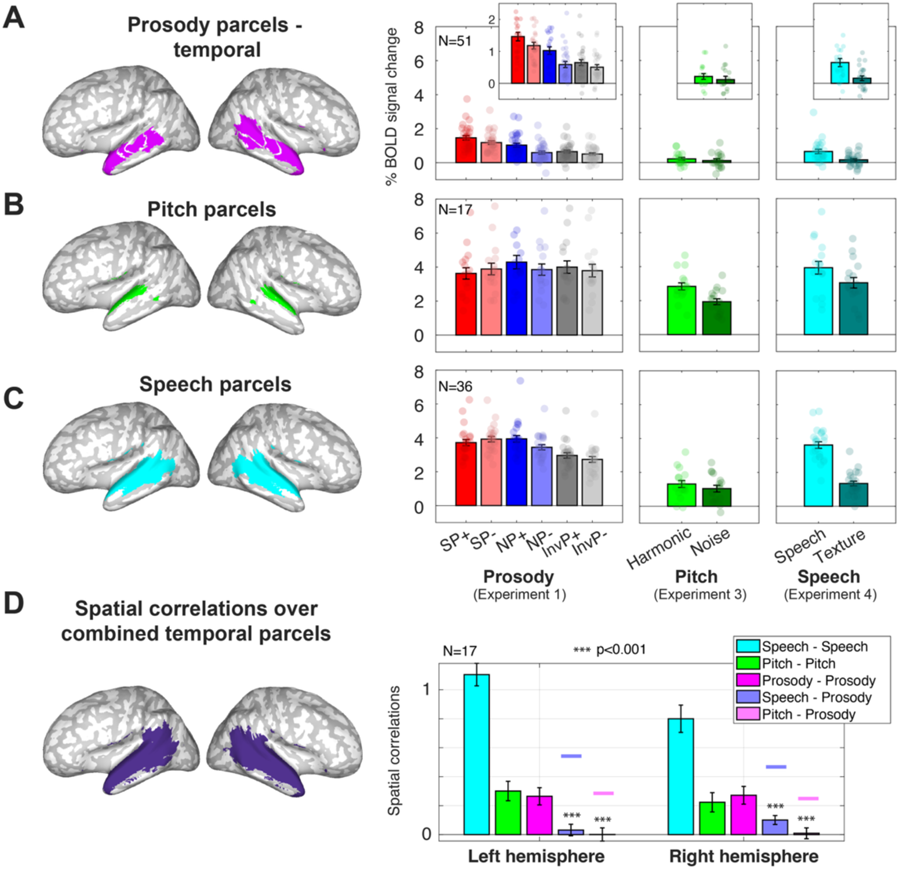
Prosody vs. auditory areas – pitch and speech (Experiments 3-4). **(A-C)** Responses in the temporal prosody areas (A), pitch areas (B), and speech areas (C) to the prosody, pitch, and speech tasks (<u>Methods</u>). We show the parcels, which are used to constrain the definition of individual-level fROIs; the fROIs are 10% of the parcels in size. Bar graphs represent the average (across participants) responses, in % BOLD signal change relative to the fixation baseline, to each condition extracted from individual-level fROIs (error bars are standard error of the mean across participants, and dots correspond to individual participants). For the prosody areas (A), responses are averaged per participant across all temporal areas (<u>Methods</u>). The inset bar graphs use a magnified y-axis scale to make the patterns more visible. (**D**) Spatial correlations (Fisher-transformed Pearson correlation coefficient) of voxel activation patterns within and between the tasks. The correlations are computed between independent subsets of the data (odd and even runs, <u>Methods</u>) and across the sets of voxels falling within a large left or right temporal lobe mask (<u>Methods</u>) presented in the inset brain.

In addition to examining the relationship with these two known auditory areas, we also examined the responses of the prosody areas to diverse natural sounds (Figure 2, Experiment 5; Boebinger et al., 2020; Norman-Haignere et al., 2013), which could be grouped into seven categories (with 2-5 clips per category; stimuli are available on OSF): English speech, foreign speech (including German, Spanish, and Italian), human non-speech vocalizations (such as yawning), human non-vocal sounds (such as walking), music, environmental sounds (such as ocean waves), and mechanical sounds (such as an electric drill). This experiment replicated—using yet another set of stimuli—the strong response of the prosody areas to English speech (characterized by natural-like prosody) and established the selectivity of this response relative to other sound categories, including music (**Figure S4**). Interestingly, foreign speech elicited a relatively low response in the prosody areas, lower than the response to nonword sentences uttered with English-like prosody, which may suggest that these areas are tuned to familiar prosody patterns of one’s native language, or that stimuli with non-native phonetics are treated as out-of-domain for these areas (see **Figure S4** and <u>Discussion</u>).

### 2.3 Prosody-responsive areas are distinct from the Multiple Demand areas, sensitive to general cognitive and attentional demands

Next, we investigated the relationship between the prosody-responsive areas and the network of brain areas that support diverse goal-directed behaviors—the so-called Multiple Demand (MD) network (Duncan, 2010; Shashidhara et al., 2019; Assem et al., 2020). The MD areas show strong modulation by cognitive difficulty (Duncan & Owen, 2000; Fedorenko et al., 2013; Hughdahl et al., 2015; Shashidhara et al., 2019; Assem et al. 2020) and attentional demands (Corbetta & Shulman, 2002; Pessoa et al., 2003). Given that prosody is often used to draw attention to particular parts of the utterance by making certain words more acoustically prominent (Terken & Hermes, 2000; Lialiou et al., 2024) some of the frontal prosody areas may simply respond to generally greater attentional demands associated with speech signals enriched with prosody (see also Kristensen et al., 2013 for evidence that certain prosodic manipulations engage general attention areas).

We used an established contrast between a harder and an easier condition of a spatial working memory task (**Figure 2**, Experiment 6; Assem et al. 2020; <u>Methods</u>) to identify the MD areas. Replicating past studies (Fedorenko et al., 2013), this contrast elicited activity in bilateral frontal and parietal areas (**Figure 4D-E**). Critically, the lateral frontal and parietal MD areas, as well as the medial-frontal MD areas responded weakly to the conditions of the prosody experiment (**Figure 4D-E**). And conversely, the prosody areas (**Figure 4A-C**) responded weakly to the spatial working memory conditions. In fact, the temporal (**Figure 4A**) and medial-frontal (**Figure 4C**) prosody areas showed the reverse pattern from the MD areas, with a stronger response during the easier condition (although both conditions fall at or below the fixation baseline). The lateral-frontal prosody areas (**Figure 4B**) showed a small Hard>Easy effect that did not reach significance (p>0.05, **Table S3**), and the overall response to these conditions was low.

The analyses of multivariate activation patterns reinforce the distinctness of the prosody areas from the MD areas: the activation pattern was more stable for the spatial working memory contrast than for the prosody contrast (**Figure 4F-H**), but critically, the activation patterns between the prosody and MD contrasts were less similar compared to the similarity of voxel activation patterns within tasks across runs (p<0.001 for all areas, LME, Methods; **Figure 4F-H, Table S6**.

**Figure 4.**
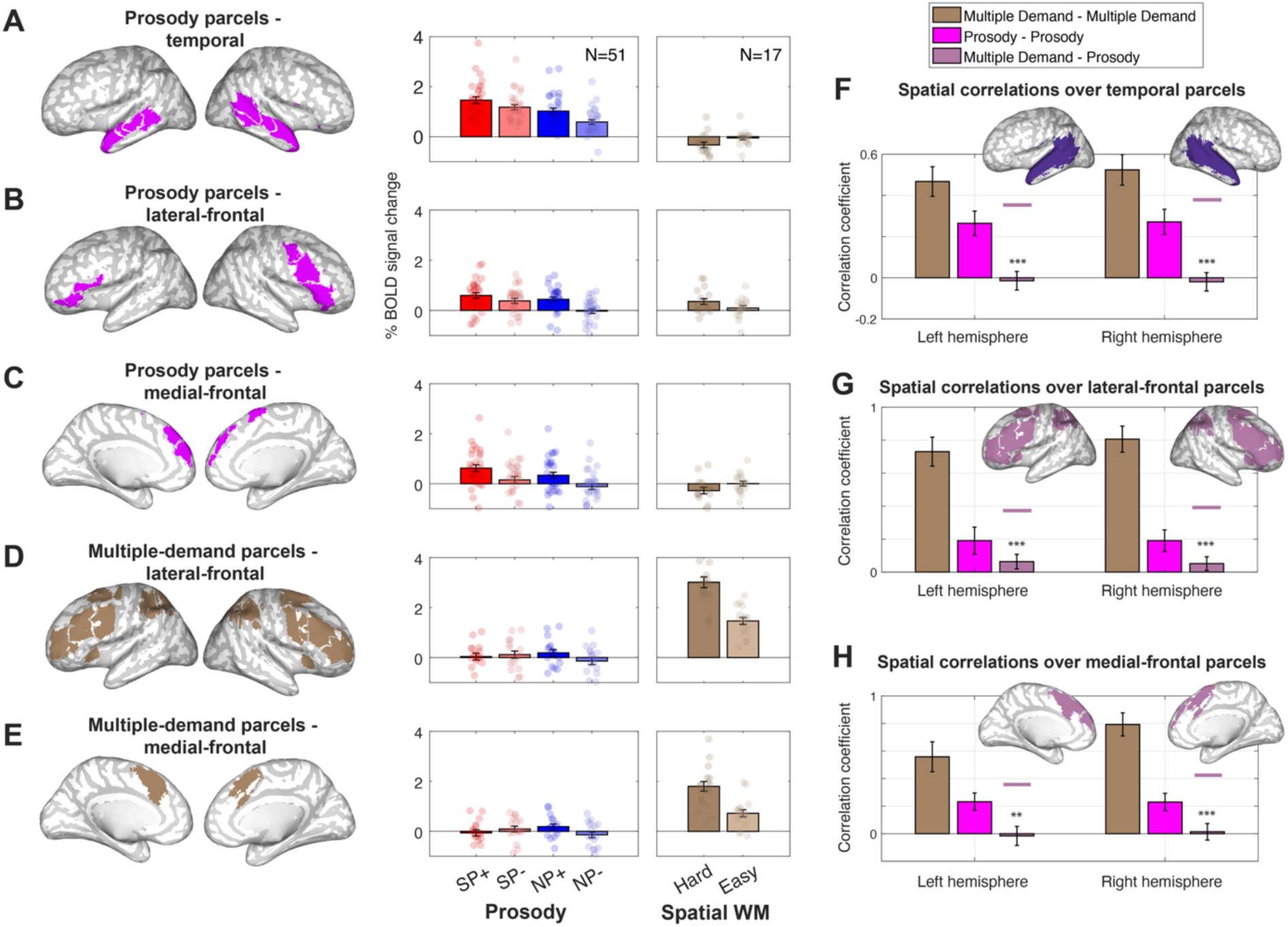
Prosody vs. Multiple Demand (MD) areas. (**A-E**) fMRI responses in the prosody and MD areas to the prosody and spatial working-memory tasks (<u>Methods</u>). (**A**) Temporal prosody areas, (**B**) lateral-frontal prosody areas, (**C**) medial-frontal prosody areas, (**D**) lateral-frontal and lateral-parietal MD areas, (**E**) medial-frontal MD areas. Bar graphs represent average across participants of % BOLD signal change relative to the baseline task of viewing a fixation cross, and error bars represent the standard error of the mean. Individual dots represent individual participants. (**F-H**) Spatial correlations (Fisher-transformed Pearson correlation coefficient) of voxel activation patterns between and within the tasks. The correlations are computed between independent parts of the data (odd or even runs, <u>Methods</u>) and across the sets of voxels falling within large left or right (**F**) temporal, (**G**) frontal-lateral or (**H**) frontal-medial lobe masks (<u>Methods</u>) presented in the inset brains.

### 2.4. Prosody-responsive areas are dissociable from, but partially overlapping with, the language areas

We next asked whether the prosody areas are distinct from left-lateralized temporal and frontal areas that specialize for language processing, often termed the *language network* (Fedorenko et al., 2024). These areas respond strongly and consistently to language—spoken, written, or signed—but show little or no response to diverse inputs and tasks, including complex auditory stimuli (such as music; Chen et al., 2023) and non-verbal communicative stimuli (such as dynamic faces and gestures; Deen et al., 2015; Pritchett et al., 2018; Jouravlev et al., 2019). These areas appear to support computations related to retrieving words from memory and combining them into structured representations (e.g., Pallier et al., 2011; Fedorenko et al., 2020; Shain, Kean et al., 2024).

The comparison of the prosody areas and the language areas is especially important given i) the role of prosody in conveying linguistic information (Snedeker & Trueswell, 2003; Watson & Gibson, 2004; Hirschberg, 2006; Breen et al., 2010; Cole, 2015; Prieto, 2015; Büring, 2016; Hellbernd & Sammler, 2016; Tomasello et al., 2022; Ladd & Arvaniti, 2023; Degano et al., 2024), ii) broad anatomical similarity between the prosody areas we identified and the language network (**Figure 5**; although it is important to note that similarity in group-level maps does not imply overlap within individual participants; e.g., Braga & Buckner, 2017; Du et al., 2024; Fedorenko et al., 2012; Fedorenko & Blank, 2020; Ladwig et al., 2025), and iii) the main effect of linguistic intelligibility that we found in the prosody areas.

To identify the language network, we used an established contrast between reading sentences and lists of nonwords (Figure 2, Experiment 7; Fedorenko et al., 2010, 2024). Although this particular localizer uses written stimuli, ample evidence exists for the modality-independence of the language network (e.g., Deniz et al., 2019; Fedorenko et al., 2010; Malik-Moraleda et al., 2022; Regev et al., 2013; Vagharchakian et al., 2012). Replicating a large body of past work (reviewed in Fedorenko et al., 2024), this contrast elicited activity in temporal and frontal areas, with stronger responses in the left hemisphere (**Figure 5C, D**). Interestingly, the responses of these language areas to the conditions of the prosody experiment resembled those of the prosody areas (**Figure 5A-D**), but with three critical differences. *First*, the language areas, especially in the left hemisphere (LH), displayed substantially greater sensitivity to verbal content compared to the prosody areas, evident in the sentences vs. nonwords contrasts from both the language localizer and the prosody experiment (p<0.001; interaction between the content effect and the type of network [prosody or language], LME, <u>Methods</u>, **Table S5**; **Figure 5A, C**). *Second*, whereas the prosody areas showed strong sensitivity to prosody in both sentences and nonword stimuli (Section 2.1), the LH language areas responded similarly strongly to sentences with and without prosody (p>0.05, LME model comparing SP+ to SP-, <u>Methods</u>, **Table S4**), presumably because the content is fully intelligible even when the prosody is disrupted (in the SP-condition). However, the language areas did show a greater response to the nonwords condition with prosody (NP+) compared to nonwords with disrupted prosody (NP-, p<0.001, <u>Methods</u>, **Table S4**), which suggests that in the absence of comprehensible content, prosody may aid in deriving some linguistic structure/meaning (see <u>Discussion</u>). And *third*, although the RH language areas’ responses to the prosody conditions were broadly similar to those of the RH prosody areas, the intelligible sentence condition with disrupted prosody (SP-) elicited a stronger response than the prosody-rich unintelligible (nonwords) condition (NP+) in the language areas, but not in the prosody areas (p<0.001; interaction between the SP-vs. NP+ effect comparing SP- and NP+ and the type of network [prosody or language], LME, <u>Methods</u>, **Table S5; Figures 5A-D**).

Furthermore, in spite of the similarity of the profiles between the prosody and the language areas (especially in the right hemisphere), the analyses of multivariate activation patterns highlighted their dissociability. For the language contrasts, we used the reading-based language localizer contrast, but also leveraged the listening-based language contrast between sentences and nonwords built into the prosody localizer (SP+,SP-> NP+,NP-). The reading-based contrast was more stable than the listening-based contrast, and both of these language contrasts were more stable than the prosody contrast (**Figure 5E**). Critically, however, the activation patterns between the prosody and either language contrast were less similar compared to the similarity of voxel activation patterns within tasks across runs (ps<0.01 for all areas except for the RH frontal regions; LME, <u>Methods</u>, **Figure 5E, Table S6**).

**Figure 5.**
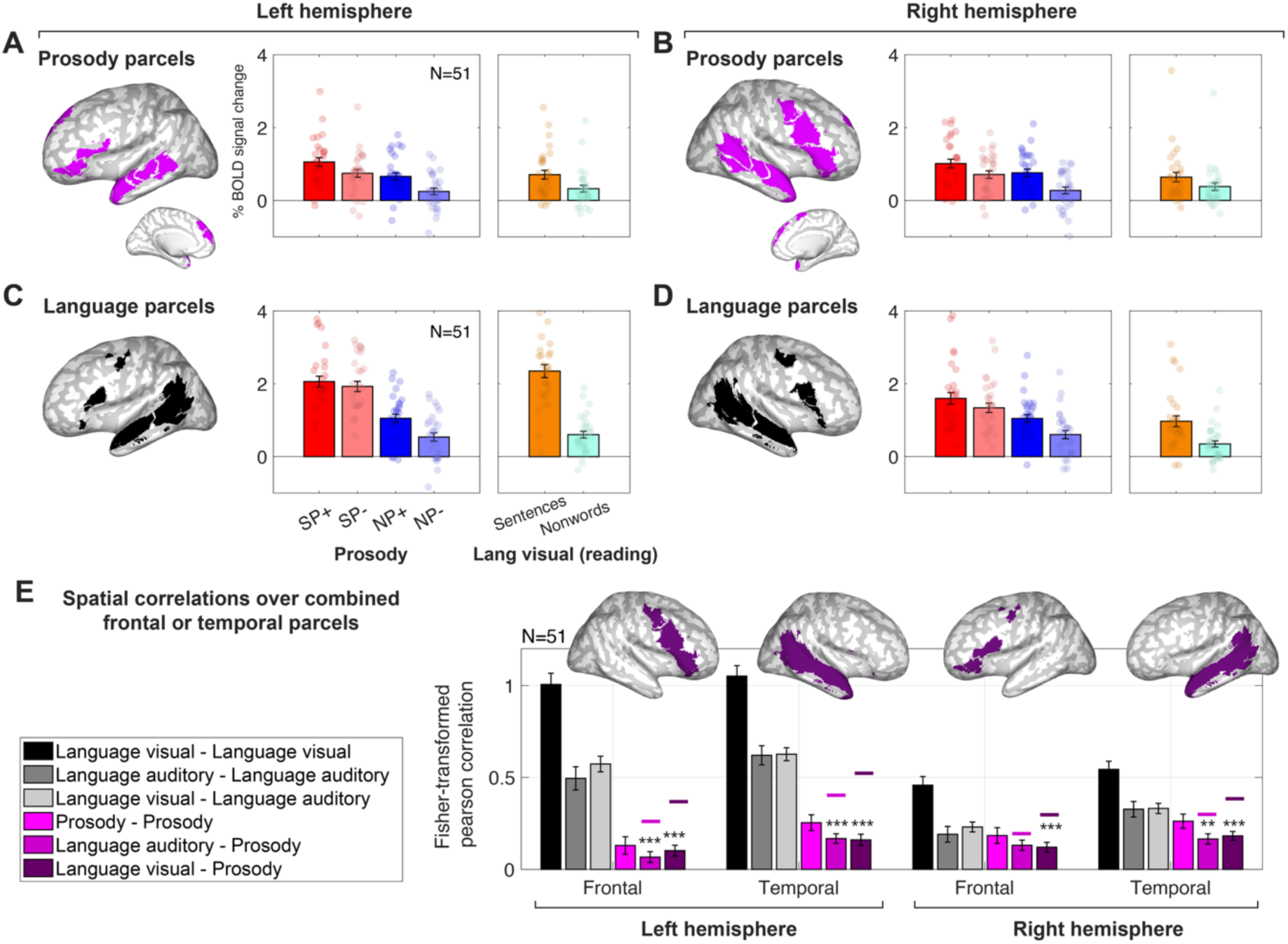
Prosody vs. language areas. (**A-D**) fMRI responses in the prosody and language areas to the prosody and (reading-based) language localizer tasks (<u>Methods</u>). (**A**) Left-hemisphere prosody areas, (**B**) right-hemisphere prosody areas, (**C**) left-hemisphere language areas, (**D**) right-hemisphere language areas. Bar graphs represent average across participants of % BOLD signal change relative to the baseline task of viewing a fixation cross, and error bars represent the standard error of the mean. Individual dots represent individual participants. (**E**) Spatial correlations (Fisher-transformed Pearson correlation coefficient) of voxel activation patterns between and within the tasks. The auditory language contrast was constructed from the prosody task data, contrasting the sentences to the nonwords, irrespective of prosody. The correlations are computed between independent parts of the data (odd or even runs, <u>Methods</u>) and across the sets of voxels falling within large left or right temporal or frontal-lateral lobe masks (<u>Methods</u>) presented in the inset brains.

### 2.5. Prosody-responsive areas are dissociable from, but partially overlapping with, the areas that process dynamic faces

Finally, we examined the relationship between the prosody-sensitive areas and areas that specialize for dynamic face perception (Pitcher et al., 2011; Pitcher & Ungerleider, 2021; Prabhakar et al., 2025). During face-to-face interactions, facial movements provide a host of communicatively-relevant signals, which—similar to prosody—can affect the interpretation of linguistic messages. To identify these face-processing areas, we used an established contrast between dynamic faces and dynamic object stimuli (Pitcher et al., 2011). Consistent with previous studies, this contrast elicited activity in bilateral superior temporal areas (**Figure 6B**). Interestingly, the responses of these face-processing areas resembled those of the temporal prosody areas (**Figure 6A**), with two differences. *First*, the face-processing areas showed substantially greater sensitivity to the faces > objects contrast compared to the prosody areas (p<0.001; interaction between the faces vs. objects effect and the type of network [prosody or dynamic faces], LME, <u>Methods</u>, **Figure 6A-B, Table S5**). And *second*, the prosody areas showed numerically greater sensitivity to the Prosody+ > Prosody-contrast compared to the face-processing areas, although the interaction was not reliable.

Similar to what we saw for the comparison of the prosody and language areas, the analyses of multivariate activation patterns highlighted the dissociability of the prosody areas and the face-processing areas. The faces > object contrast was more stable than the prosody contrast (**Figure 6C**), but critically, the activation patterns were more similar across runs within the prosody contrast compared to between the prosody and face processing contrasts bilaterally (*p*s<0.05; <u>Methods</u>, **Figure 6C, Table S6**).

**Figure 6.**
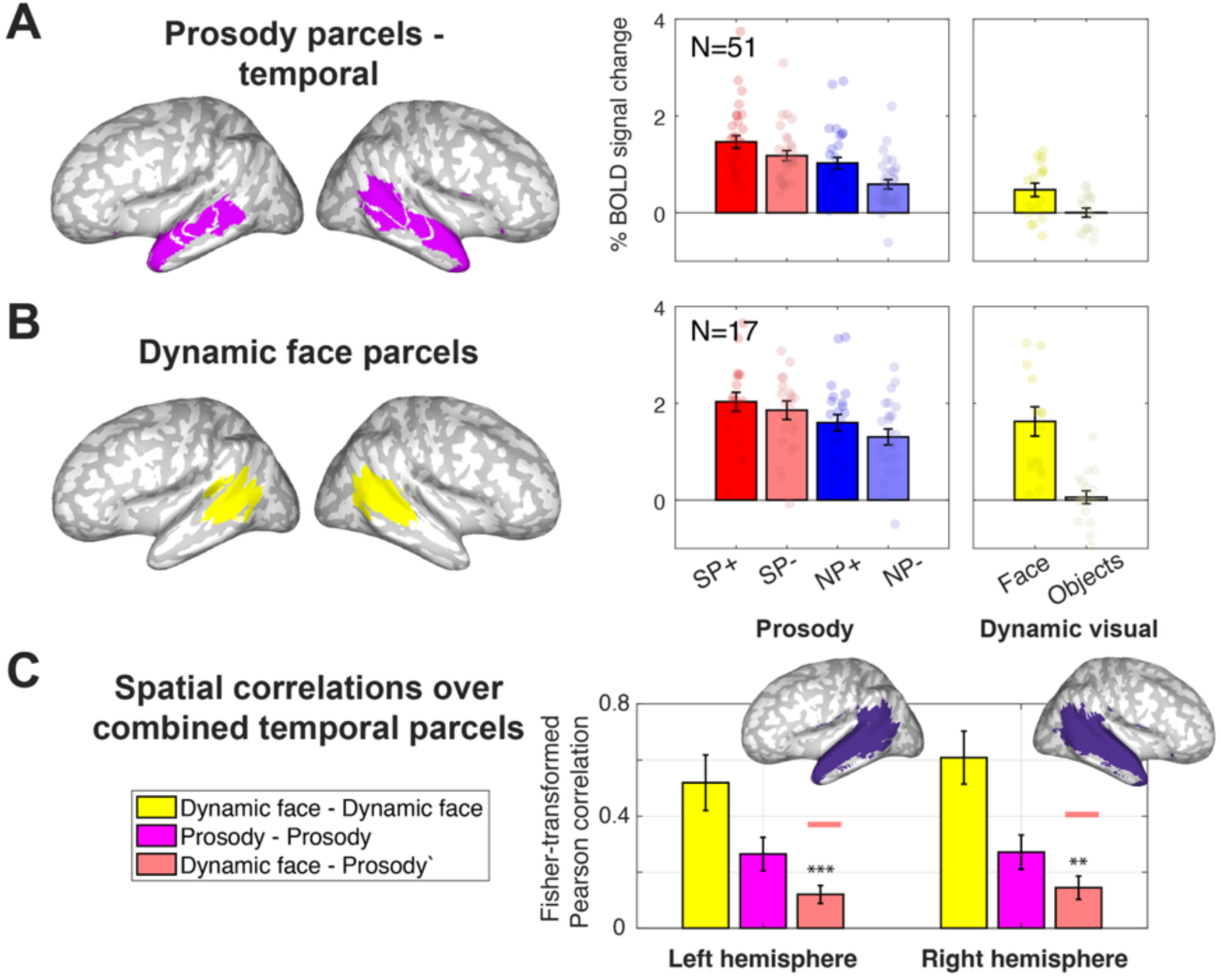
Prosody vs. dynamic face areas. (**A-B**) fMRI responses in the prosody and dynamic face areas to the prosody and dynamic face perception tasks (<u>Methods</u>). (**A**) Temporal prosody areas, (**B**) temporal dynamic face areas. Bar graphs represent average across participants of % BOLD signal change relative to the baseline task of viewing a fixation cross, and error bars represent the standard error of the mean. Individual dots represent individual participants. (**C**) Spatial correlations (Fisher-transformed Pearson correlation coefficient) of voxel activation patterns between and within the tasks. The correlations are computed between independent parts of the data (odd or even runs, <u>Methods</u>) and across the sets of voxels falling within large left or right temporal lobe masks (<u>Methods</u>) presented in the inset brains.

## 3. DISCUSSION

Using a novel fMRI paradigm, we identified a set of bilateral temporal and frontal brain areas that are robustly sensitive to prosody whether with or without comprehensible verbal content. The temporal areas showed generalizable responses to new prosody-rich stimuli and were distinct from auditory areas that specialize for the processing of pitch and speech sounds, as well as from areas sensitive to general attentional demands. They were also dissociable from the language areas and from areas that process dynamic faces—even in the right hemisphere—but also showed some degree of overlap with both. The frontal areas showed poorer generalizability to new stimuli but were clearly distinct from areas sensitive to attentional demands and partially dissociable from the frontal language areas. Together, these findings suggest that the core areas that processes speech prosody are located in the bilateral temporal lobes and occupy a distinct functional niche within the brain’s architecture, but also show some degree of integration with the language areas and dynamic-face perception areas (**Figure 7**). In the following sections, we discuss these findings in relation to the current theoretical and empirical landscape of research on human communication.

**Figure 7.**
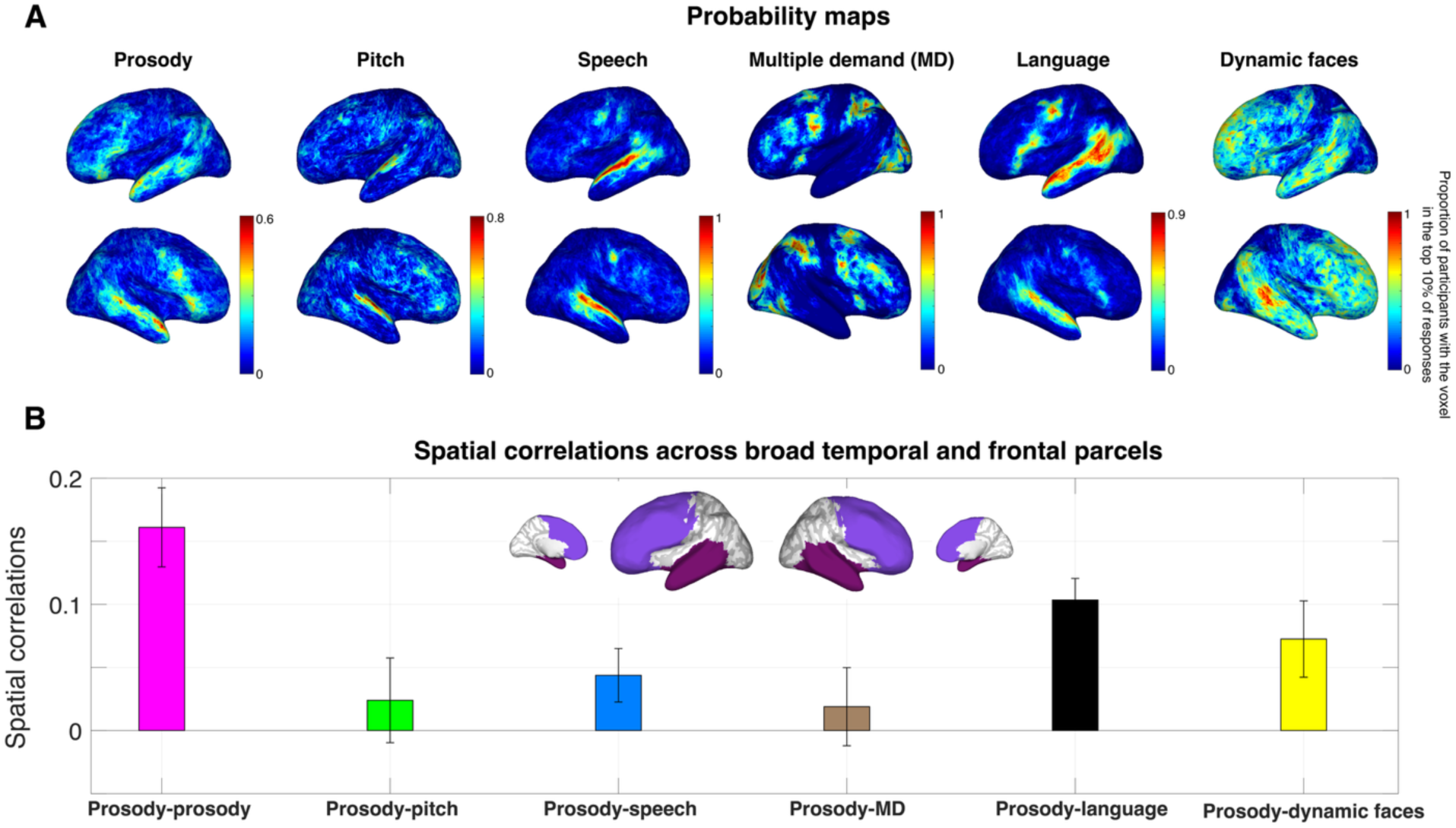
A summary of the relationship between the prosody areas and all other tested areas: Pitch, speech, multiple demand, language, and dynamic face processing areas. (**A**) Probabilistic activation overlap map for each contrast: prosody (as in Figure 1E, N=51), pitch (N=17), speech (N=37), multiple demand (N=17), language (N=51) and dynamic faces (N=17) (<u>Methods</u>). (**B**) Spatial correlations (Fisher-transformed Pearson correlation coefficient) of voxel activation patterns between and within the tasks. The correlations are computed between independent parts of the data (odd or even runs, <u>Methods</u>) and across the sets of voxels falling within large bilateral temporal and frontal lobe masks (<u>Methods</u>) presented in the inset brains. The correlations were first computed for the temporal lobe and for the frontal lobe separately, and then averaged across the lobes per participant. Bar graphs represent the mean across participants, and error bars represent the standard error of the mean.

### 3.1. The timescale of prosody

Prosody relies on auditory features, including pitch, loudness, and duration, which are primarily processed in the auditory cortex. These auditory brain areas are engaged by both simple stimuli, such as pure or harmonic tones (Petkov et al., 2006; Taaseh et al., 2011; Norman-Haignere, Kanwisher, & McDermott, 2013), and by complex prosody-rich speech stimuli (Engineer et al., 2008; Rauschecker & Scott, 2009; Tang et al., 2017). Crucially, however, prosody is not defined just by isolated acoustic cues but by how they unfold over time—while local pitch movements may span only one–two syllables (100–500 ms), larger prosodic units extend across about a second or more (Inbar et al., 2020, 2025), and prosodic interpretation strongly depends on longer linguistic context spanning multiple words (Cole, 2015; Regev et al., 2025). To isolate the processing of these temporally-extended prosodic patterns, we contrasted speech stimuli with natural prosody vs. stimuli whose prosody has been disrupted by splicing together words recorded in isolation. This contrast engaged temporal regions outside the auditory cortex, along the superior temporal sulcus (STS), as well as lateral and medial frontal regions (for consistent past findings, see Fedorenko et al., 2013).

A plausible reason for why prosody-sensitive areas showed no overlap with either the auditory areas specialized for processing pitched sounds, or the auditory speech-processing areas, is that the temporal integration window of those auditory areas is relatively short. The temporal integration window of the speech-processing areas is up to approximately half a second (Overath, McDermott, et al., 2015; Norman-Haignere et al., 2022). For the pitch areas, the integration window appears to be even shorter, up to about 200 ms (Norman-Haignere, Kanwisher, & McDermott, 2013; Norman-Haignere et al., 2022). In contrast to these auditory areas, the language areas integrate information over a longer timescale, of between 1 and 8 words (Blank & Fedorenko, 2020; Caucheteux et al., 2023; Jain et al., 2020; Lerner et al., 2011; Regev et al., 2024; Shain et al., 2024). And although the language and the prosody areas could be dissociated in the current study, their functional profiles and activation topographies exhibited some similarity. Comparable timescales of information processing may partially underlie this similarity, although the timescale of prosodic processing requires further investigation.

### 3.2. Prosody may constitute a special type of a non-verbal communicative signal

Much past work on human communication has focused on the verbal component of language. However, recent years have witnessed a sharp increase in research on the processing of non-verbal communicative signals that accompany speech and a growing emphasis on the multimodal nature of naturalistic human interactions (Goldin-Meadow, 1999, 2014; Hirschberg, 2002; Vigliocco et al., 2014; Holler & Levinson, 2019; Hadley et al., 2022; Hagoort & Özyürek, 2024). Indeed, prior to the invention of writing and other technologies that allow us to carry out linguistic exchanges without the full multi-modal experience, human communication consisted primarily of face-to-face interactions, which require integrating linguistic information with all the other cues a speaker produces: from eye gaze and facial expressions, to gestures and body language, to prosody. And these non-verbal cues are often critical to correctly interpreting the utterance and understanding speaker intent (Archer & Akert, 1977; Thompson & Massaro, 1986; Wharton, 2009).

And yet, although human communication is often multimodal, it need not necessarily be. We can talk to each other in complete darkness and across visual obstacles, removing all visual signals. Besides, many visual and auditory signals—such as a smile, or eyes widening, or a sigh—are only present occasionally during linguistic exchanges. Perhaps the fact that non-verbal communicative signals do not always or continuously accompany linguistic interactions leads to the separation of their underlying neural mechanisms. Indeed, the brain areas that process language show little or no response to facial expressions, gestures, and vocalizations, such as sighs and laughter (Deen et al., 2015; Pritchett et al., 2018; Jouravlev et al., 2019). Instead, these non-verbal communicative signals are processed by distinct visual and auditory brain areas, ruling out the idea that a single brain system supports the processing of all communicative information.

However, prosody differs from other non-verbal communication channels in that it *always* accompanies speech, being inseparable from the speech signal. This inseparability holds even in whispered speech (Heeren & van Heuven, 2014) and when processing written language—a phenomenon known as *implicit prosody* (Breen, 2014); and prosody is also present in sign languages (Sandler, 2010; Brentari et al., 2018; Sandler et al., 2020). In our study, we found that although the fine-grained activation topographies are distinct between language processing and prosody processing, and the functional profiles are not identical, they show some similarity, especially in the right temporal lobe. This partial integration between the system that processes verbal content, and the system that processes prosodic information aligns with recent work, which has found a substantial degree of redundancy in the information carried by these two communication channels: words that are particularly informative (as can be measured using information-theoretic approaches; Shannon, 1948) are also marked prosodically, being uttered louder, slower, and with higher pitch peaks (Seyfarth, 2014; K. Tang & Shaw, 2021; Wolf et al., 2023; Clark et al., 2025; Regev et al., 2025). Moreover, next word prediction is more accurate when human listeners (and AI models) have access to the full speech signal – which includes prosody compared to the text alone (Botch & Finn, 2025). Thus, prosodic signals appear to complement and reinforce linguistic messages in the service of efficient communication.

Interestingly, we also found some overlap between the brain areas that process prosody and those that process dynamic faces. The fact that prosodic processing overlaps with both the language areas and the areas that process visual communicative information may suggest that it provides a critical interface between these two types of communication and perhaps serves an integratory role. However, whether prosody areas overlap with brain areas that process other non-verbal communicative signals, and how exactly the orchestration of different communicative signals happens during face-to-face communication remains to be discovered.

### 3.3. The function of the prosody areas and the nature of prosodic representations

Given the response profile of the prosody-processing brain areas, what can we infer about their function? Can we make some guesses based on what we know about the language areas, which we found to overlap with the prosody areas? The language areas support both comprehension and production (Menenti et al., 2011; Silbert et al., 2014; Hu et al., 2023), suggesting that they store mappings between form and meanings, needed both for extracting meaning from word sequences (comprehension) and converting thoughts into verbal utterances (production) (Fedorenko et al., 2024). By analogy, prosody-responsive areas may learn, store, and use mappings between sets of acoustic cues, or even specific contours, and meanings, and then use these when generating or interpreting prosodic contours.

The nature of prosodic representations, how they correspond to particular meanings, and more generally, what kinds of meanings prosody can convey remain an active area of research (Hirschberg, 2006; Prieto, 2015; Westera et al., 2020). In contrast to the words themselves, which refer to objects, actions, and events/states, prosodic contours convey pragmatic, discourse-related, and often less clearly categorical distinctions, such that phrases and sentences with very different lexical content can express the same prosodically marked meaning. For example, prosody can signal whether an utterance is a statement, a question, or a request (Hirschberg, 2002; Gunlogson, 2004; Bartels, 2014; Hellbernd & Sammler, 2016), or mark the ‘information structure’ of a sentence (Jackendoff, 1972; Féry & Krifka, 2008), including what information is assumed to be known by the interlocutors, what information is new, what information is particularly important, etc. (Lieberman, 1960; Bolinger, 1961; Cooper et al., 1985; Beckman, 1986; Ito et al., 2004; Breen et al., 2010; Katz & Selkirk, 2011; Chodroff & Cole, 2019; Roettger et al., 2019; see Cole, 2026 for a review). Prosodic contours can also signal the speaker’s affect, or their social status (Banse & Scherer, 1996; Laukka et al., 2005; for a review see Larrouy-Maestri et al., 2025).

How these different meanings correspond to the acoustic signal has been a matter of debate. Some approaches map acoustic features, such as pitch or loudness, directly onto meaning categories (Lieberman, 1960; Cooper et al., 1985; Bartels & Kingston, 1994; Pell, 2001; Xu & Xu, 2005; Breen et al., 2010; Cole et al., 2010), whereas others first reduce the dimensionality of speech acoustics into a small set of prosodic categories, which are then linked to meanings (the *intonational phonology* approach; Pierrehumbert, 1980; Gussenhoven, 1983; Silverman et al., 1992; Dilley & Brown, 2005; Beckman et al., 2005; Ladd, 2008). One limitation of both approaches is that they assume a small number of *a priori* meaning categories, and intonational phonology further requires extensive manual annotation with only moderate inter-rater agreement (Breen et al., 2012; cf. Cole & Shattuck-Hufnagel, 2016). An alternative more recent approach instead attempts to discover types of prosodic patterns directly from large-scale speech corpora, without assuming a predefined inventory of either prosodic forms or meaning categories (e.g., Matalon et al., 2025). This bottom-up approach faces its own challenges, such as understanding the meanings encoded by the discovered prosodic patterns, including whether they can be grouped into fewer functionally coherent categories, and how they combine into larger prosodic structures.

In summary, given the topographic similarity and partial overlap between the prosody network and the language network, a reasonable hypothesis may be that the prosody areas store learned prosodic knowledge representations, similar to how the language areas store verbal-linguistic representations. The fact that the prosody areas responded more strongly to nonword sequences with English-like prosody than to foreign speech further suggests that these areas may be attuned to the prosodic patterns of one’s native language. Such tuning, which may start as soon as hearing develops in utero (Gervain, 2018), may allow listeners to extract some information/meaning even from meaningless sequences, which may be important both for language learning and speech processing under noisy conditions. Alternatively, foreign speech with unfamiliar phonology may be treated as out-of-domain for these brain areas. Testing these possibilities will require stimuli that cross familiar versus unfamiliar prosody with familiar versus unfamiliar phonology, or foreign languages that vary in their prosodic similarity to English. More broadly, further evaluating the representational nature of the prosody areas will benefit from probing these areas’ engagement during prosody *production* as well as comprehension. And going forward, neuroscientific investigations of prosodic processing should synergize with ongoing work in linguistics and cognitive science aiming to characterize linguistic representations and understand the way in which they are used during communication.

### 3.4. Limitations and open questions

This study lays a foundation for future investigations of prosody-responsive brain areas, but it leaves many questions about prosodic representations and computations open. First, by design, we targeted prosody *broadly*, without trying to examine its different hypothesized functions. Our disrupted-prosody condition impaired both pitch and timing cues, preventing the emergence of phrasal-level contours and removing pragmatic and emotional/social meaning cues. This choice was primarily driven by a methodological consideration: we wanted to initially cast a wide net to include all prosody-relevant brain areas. In addition, the same acoustic cues—pitch, loudness, and duration/tempo—contribute to conveying different kinds of meanings, albeit in different ways (Turner, 2025), and in a large-scale behavioral investigation of individual differences in non-literal comprehension abilities, linguistic and emotional prosody tasks loaded on the same latent factor (Floyd, Jouravlev et al., 2025). However, past studies of patients with brain damage have emphasized the greater importance of the right hemisphere for emotional prosody, and of the left hemisphere—for linguistic prosody (Witteman et al., 2011; Weed & Fusaroli, 2020). Future examination of responses in the prosody areas to diverse linguistic and emotional contrasts will be necessary to test for this putative hemispheric dissociation, and to search for other dissociations among the different prosody-responsive areas.

Second, we have focused on *English*, which is not representative of many of the world’s languages (Blasi et al., 2022). Languages vary in how they use prosody (Hirst & Di Cristo, 2001; Jun, 2006; Lee et al., 2015) and so-called tonal languages, such as Mandarin and Somali, use pitch cues not only for phrasal prosody, but also to mark lexical or grammatical distinctions (Bolinger, 1978; Hyman, 2016). Whether the brain areas discovered here support prosody perception in speakers of diverse languages, including sign languages, remains to be discovered.

Third, we have examined mature brains. Are these prosody areas already present in young infants’ brains, given that prosody provides a critical scaffold for linguistic development (Shukla et al., 2011; Gervain & Werker, 2013; Graf Estes & Hurley, 2013; Piazza et al., 2017), or even at birth, given that prosodic information can be heard in the womb (DeCasper & Spence, 1986; Moon et al., 1993; Nazzi et al., 1998; Gervain, 2018; Martinez-Alvarez et al., 2022)? These areas may be even more spatially extensive in infants than in adults, encroaching on the areas that eventually become language-selective. Developmental studies can investigate this possibility.

Fourth, we have found overlap between the prosody areas and areas that process dynamic faces. Do prosody areas also overlap with other areas that process non-verbal communicative signals, such as voices (Belin, Zatorre, Lafaille, et al., 2000) or body language/gestures (Jouravlev et al., 2019), or others’ social interactions (Isik, Koldewyn et al., 2017)? If prosody serves as an interface between linguistic and non-verbal communication, it may show some degree of overlap with all these other communication-relevant areas.

Finally, we have used precision fMRI, which is the best available approach for identifying cortical areas supporting particular functions of interest, but is not well-suited for understanding the dynamics of information processing or inter-system interactions. Future work can use this localizer contrast in studies that use intracranial approaches to probe the timecourse of prosodic processing, and investigate fine-grained temporally-locked distinctions in both linguistic and emotional prosody. Multivariate fMRI approaches and approaches that compare neural responses with representations extracted from neural network models (similar to what has been done for vision and language; Yamins & DiCarlo, 2016; Kell & McDermott, 2019; Richards et al., 2019; Toneva & Wehbe, 2019) can also be useful to further characterize the information represented and computed in the prosody areas.

In conclusion, we have identified a set of brain areas that respond robustly to prosody, are distinct from known auditory and general attention areas, but show partial overlap with the language areas and areas that process dynamic faces. These areas can now be studied in even greater detail in future work, to continue uncovering the complex orchestration of many distinct components of human communication.

## 4. METHODS

### 4.1 Participants

In total, 51 individuals (age 18-46; mean 24.8 +/-7 years; 29 (56.8%) females) from the Cambridge/Boston, MA community participated in fMRI experiments for payment. Forty-eight of the 51 participants were right-handed (∼94%), as determined by the Edinburgh handedness inventory (Oldfield, 1971), or self-report; the remaining participants were left-handed (n=3; see Willems et al., 2014 for arguments for including left-handers in cognitive neuroscience research).

All participants were native English speakers, and all provided written informed consent in accordance with the requirements of MIT’s Committee on the Use of Humans as Experimental Subjects (COUHES), which approved the study.

**Table 1.** Details of the dataset. 51 participants performed the prosody localizer experiment and different subsets of the other experiments. 7 of the participants completed experiments across two or three scanning sessions.

| N | Exp 1<br>Prosody<br>localizer | Exp 2<br>Generalization<br>to new stimuli | Exp 3<br>Pitch<br>localizer | Exp 4<br>Speech<br>localizer | Exp 5<br>Diverse<br>natural<br>sounds | Exp 6<br>MD<br>network<br>localizer | Exp 7<br>Language<br>localizer | Exp 8<br>Dynamic face<br>localizer |
| --- | --- | --- | --- | --- | --- | --- | --- | --- |
| 19 | ☒ | ☒ | ☐ | ☐ | ☐ | ☐ | ☒ | ☐ |
| 12 | ☒ | ☐ | ☐ | ☐ | ☒ | ☐ | ☒ | ☐ |
| 11 | ☒ | ☐ | ☒ | ☒ | ☐ | ☒ | ☒ | ☒ |
| 3 | ☒ | ☒ in a different<br>session | ☒ | ☒ | ☒ in a<br>different<br>session | ☒ | ☒ | ☒ |
| 2 | ☒ | ☐ | ☐ | ☐ | ☐ | ☐ | ☒ | ☐ |
| 2 | ☒ | ☐ | ☒ | ☒ | ☒ in a<br>different<br>session | ☒ | ☒ | ☒ |
| 1 | ☒ | ☒ in a different<br>session | ☒ | ☒ | ☐ | ☒ | ☒ | ☒ |
| 1 | ☒ | ☒ in a different<br>session | ☐ | ☐ | ☒ | ☐ | ☒ | ☐ |
| Total<br>N | 51 | 24 | 17 | 17 | 18 | 17 | 51 | 17 |

### 4.2 Design, materials, and procedure

Each participant completed a prosody ‘localizer’ task (detailed below) and 2-7 additional tasks, which were included to characterize prosody-sensitive areas and understand their relationship to known brain systems (see **Table 1** for details). The scanning sessions lasted approximately 2 hours.

#### Experiment 1 - Prosody localizer

This task was designed to identify brain areas that support prosody perception. Participants listened to sentences or nonword lists produced with natural or list-like prosody, leading to a 2x2 blocked design with the following conditions (**Figure 1**): Sentences Prosody+ (SP+), Sentences Prosody-(SP-), Nonwords Prosody+ (NP+), and Nonwords Prosody- (NP-). Two additional conditions were constructed to control for low-level auditory differences between the Prosody+ and Prosody-conditions: Control Prosody+ (CP+) and Control Prosody- (CP-) were created by playing the stimuli in the NP+ and NP-conditions backwards (see <u>Stimulus construction</u> below for details). A trial consisted of a recording of a sentence, a nonword sentence, or reversed speech and lasted 5 seconds (the original recordings varied in duration between 2.2 s and 4.8 s [avg=3.5 s, std =0.6 s], and were padded with silence to reach 5 s). A block consisted of three trials and a silent period of 1 s where a hand icon appeared, either between the 2^nd^ and 3^rd^ trials, or after the 3^rd^ trial, for a total block duration of 16 s. Participants were instructed to press a button when the hand icon appeared, to maintain alertness. Each run consisted of 12 experimental blocks (2 for each of the 6 condition) and 3 fixation blocks (14 s each), for a total duration of 234 s (3 min 54 s). The order of conditions varied across runs and participants, but was always palindromic within a run. Each participant performed 6 runs, for a total of 12 blocks per experimental condition.

<u>Stimulus construction</u>: We extracted sentences from the transcripts of two spoken American English corpora: Switchboard (Godfrey et al., 1992) and the Corpus of Contemporary American English (COCA; Davies, 2010). We searched these corpora for sentences which included between 10 and 14 words. The word counts included disfluencies, such as “uh” and “um” (but not word repetitions). We also excluded sentences that contained strongly emotional or political content, or were confusing out of context. From the full set of 24,727 (Switchboard) and 594,737 (COCA) sentences of the target length, we identified 126 sentences (63 from each corpus) that we judged could be produced with expressive prosody. In particular, to ensure variation in both prosodic prominence and temporal cues, we wanted to include both statements and questions, sentences that contain words that could be emphasized relative to their context, and sentences consisting of phrases or clauses that could be separated by a prosodic phrase boundary. We annotated the sentences to mark the words to emphasize and the boundary locations, and had an actress record each sentence using expressive prosody. The recorded sentences were trimmed (so as to start at the onset of speech and end at the speech offset), and their durations were computed. Then, two of the authors (TIR and EF) rated each recording for naturalness on a scale from 1 to 5. Finally, we selected a subset of 72 sentence recordings that were rated as natural-sounding and that did not exceed 5 seconds in duration. These recordings served as the Sentence Prosody+ (SP+) condition.

To create the Sentence Prosody- (SP-) condition, we extracted all the individual words that comprised the 72 sentences (a total of 491 unique words), scrambled their order, and had the actress record each word in isolation with neutral, falling prosody. This typical single word prosody can be characterized as having a very modestly scaled high pitch accent marking the word-level stress (H* in the ToBI system; (Silverman et al., 1992; M. Beckman & Ayers G, 1997)followed by a modestly scaled fall to a low boundary tone at the end of the word. We then spliced together the recordings of individual words to recreate the original sentences. Prior to splicing, 30 ms linear ramps from (to) zero were added at the beginning (ending) of each recording of an individual word in order to prevent acoustic edge artifacts. These constructed recordings sounded like a disconnected list of words, where each word forms its own intonational phrase (see **Figure 1A**). Because the stimuli in the SP-condition were generally longer than the sentences in the SP+ condition, we time-compressed the SP-stimuli using the MATLAB function *stretchAudio.m* to match their corresponding sentences.

To create the Nonword Prosody+ (NP+) and the Nonword Prosody-(NP-) conditions, we replaced each word with a matched (in the number and type of syllables) pronounceable nonword using the Wuggy software (Keuleers & Brysbaert, 2010). Then, for the NP+ condition, the actress recorded the nonword sentences using prosody that matched the prosody of the corresponding sentences as closely as possible. For the NP-condition, we followed the same procedure as for the SP-, using the nonword sentences. To create the control conditions, InvP+ and InvP-, the recordings of the NP+ and NP-stimuli were played backwards, respectively. All stimuli were equalized for rms and during the scanning session, the volume was subjectively adjusted for each participant to make sure the recordings were clearly audible.

These 72 stimuli in each condition were distributed across two lists. List 1 consisted of stimuli 1-36 for the SP+, NP-, and InvP+ conditions, and stimuli 37-72 for the SP-, NP+, and InvP-conditions; and List 2 consisted of stimuli 1-36 for the SP-, NP+, and InvP-conditions, and stimuli 37-72 for the SP+, NP-, and InvP+ conditions. In this way, each list contained only a single instance of a particular sentence and a single instance of a particular prosodic contour. Approximately half of the participants were assigned to each list.

#### Experiment 2 – Generalization of the prosody results from Experiment 1

This task was designed to evaluate the generalizability of the prosody localizer results to a different set of experimental stimuli. This experiment included conditions that were conceptually similar to three critical conditions of Experiment 1: SP+, NP+, and InvP+, along with several additional conditions of no interest. SP+ and InvP+ were part of the same run types, whereas NP+ was taken from different experimental runs. We start by describing the runs containing the SP+ and InvP+ conditions (along with other conditions of no interest). In the SP+ condition, participants listened to excerpts from unscripted telephone conversations from the CALLFRIEND American English-Non-Southern Dialect Second Edition corpus. Seventeen clips with female voices (6 unique speakers) and 19 clips of male voice (7 unique speakers) were selected, with length ranging between 2.5 and 7.3 seconds. In the InvP+ condition, participants listened to the same clips played in reverse. A block consisted of three trials from the same condition and a silent period of 1 s where a hand icon appeared, for a total block duration of 16 s. The original recordings were grouped into blocks so as to have three stimuli that together last close to 15 s; shorter stimuli were padded with silence to maintain a consistent block length of 15 s. Participants were instructed to press a button when the hand icon appeared, to maintain alertness. Each run consisted of 16 experimental blocks (2 per condition) and 3 fixation blocks (12 s each), for a total duration of 228 seconds (3 min 48 s). Each participant performed 6 runs, for a total of 12 blocks per experimental condition. We next describe the runs containing the NP+ condition (along with other conditions of no interest). In the NP+ condition, participants listened to sentences where real words were replaced by nonwords and recorded by four different actors (two males, stimuli from Deen et al. 2020). A block consisted of three trials from the same condition and a silent period of 1 s where a hand icon appeared, for a total block duration of 18 s. The original recordings were grouped into blocks to have three stimuli that together last close to 17 s; shorter stimuli were padded with silence to maintain a consistent block length of 17 s. Participants were instructed to press a button when the hand icon appeared, to maintain alertness. Each run consisted of 9 experimental blocks (3 per condition) and 4 fixation blocks (12 s each), for a total duration of 210 seconds (3 min 30 s). Each participant performed 3 runs, for a total of 9 blocks per experimental condition.

#### Experiment 3 – Pitch localizer

This task was designed to target brain areas that specialize for perceiving pitched sounds, as reported in Norman-Haignere et. al. (2013, stimuli downloaded from https://web.mit.edu/svnh/ www/Resolvability/Efficient_Pitch_Localizer.html). Participants listened to sequences of harmonic tones (H) or frequency-matched gaussian noise (N) stimuli in a blocked design. The harmonic tones were designed to elicit a clear pitch percept. All tones had the same harmonic structure, consisting of harmonics 3-6 (which are resolved harmonics), but each tone had a different, constant fundamental frequency, ranging between 249 and 445 Hz (mean of 333 Hz). The frequency-matched noise stimuli were created by band-pass filtering gaussian noise at a frequency range that matched each of the harmonic tones (the range corresponding to harmonics 3-6). Critically, these stimuli elicit no clear pitch percept. A tone / noise sequence (composed of 6, 8, 10 or 12 tones / noise stimuli of equal duration) lasted 2 s and was preceded and followed by 200 ms of silence, for a total duration of 2.4 s. A trial consisted of such stimuli, with additional 100 ms long silence periods added before and after the first stimulus, for a total trial duration of 5s. A block consisted of three trials and a silent period of 1 s where a hand icon appeared, either between the 2^nd^ and 3^rd^ or after the 3^rd^ trial, for a total block duration of 16 s. Participants were instructed to press a button when the hand icon appeared, to maintain alertness. Each run consisted of 16 experimental blocks (8 per condition) and 5 fixation blocks (14 s each), for a total duration of 326 s (5 min 26 s). The order of conditions varied across runs and participants, but was always palindromic within a run. Each participant performed 2 runs, for a total of 16 blocks per experimental condition.

#### Experiment 4 – Speech localizer

This task was designed to target brain areas that specialize for perceiving speech sounds, regardless of their linguistic content, as in Overath, McDermott et al. (2015). Participants listened to nonword sentences (N) or matched control stimuli (C) in a blocked design. The nonword sentences were designed to match the phonology of English sentences (not used in this experiment) as much as possible, while lacking meaning. A female speaker (different from the speaker who recorded the prosody localizer stimuli) recorded the nonword sentences with as natural prosody as possible; she was instructed to utter each invented nonword sentence freely and naturally. The control stimuli were created using an algorithm from McDermott & Simoncelli (2011) that estimates summary statistics (averages of acoustic measures across time) from a sound file and synthesizes ‘sound textures’ via noise modulations, imposing the measured statistics. Thus, the low-level auditory features are closely matched between the two conditions, but higher-level features, which give speech its characteristic percept, differ. A trial consisted of a recording of a nonword sentence or the control stimulus and lasted 4-5 seconds (the recordings that were shorter than 5 s were padded with silence). A block consisted of 3 trials and a silent period of 1 s where a hand icon appeared, either between the 2^nd^ and 3^rd^ or after the 3^rd^ trial, for a total duration of 16 s. Participants were instructed to press a button when the hand icon appeared, to maintain alertness. Each run consisted of 16 experimental blocks (8 per condition) and 5 fixation blocks (14 s each), for a total duration of 326 s (5 min 26 s). The order of conditions varied across runs and participants, but was always palindromic within a run. Each participant performed 2 runs, for a total of 16 blocks per experimental condition.

#### Experiment 5 – Responses to natural sounds

This task is a shortened version of the task used in Norman-Haignere et al. (2015) designed to measure brain responses to a diverse set of natural sounds. This shorter version included 30 (of the original set of 165) 2-second natural sound stimuli, which spanned 7 broad categories: English speech (2 stim), Foreign speech (3 stim), Human vocal sounds (4 stim), Human non-vocal sounds (4 stim), Music (5 stim), Environmental sounds (4 stim), and Mechanical sounds (5 stim); 3 other stimuli were not assigned to a particular category. Stimuli were presented in a “mini-block design,” where each 2-second sound was repeated three times in a row. Each stimulus was presented in silence, with a single fMRI volume collected between each repetition (i.e., “sparse scanning”, Hall et al., 1999). Either the second or the third repetition in each mini-block was 8 dB quieter, and participants were instructed to press a button when they heard the quieter sound, to maintain alertness. Each run consisted of 30 mini-blocks, one for each stimulus, for a total duration of 408 s (6 min 48 s). Each participant performed 6 runs for a total of 6 mini-blocks per stimulus.

#### Experiment 6 – Multiple demand network localizer

This task was designed to target brain areas that are sensitive to general cognitive demands associated with increased attention, working memory, and related executive processes; the network of brain areas that support these processes is known as the multiple demand (MD) network (Duncan, 2010; Duncan et al., 2020). Participants performed a spatial working memory task consisting of a Hard (H) and an Easy (E) condition in a blocked design; the Hard>Easy contrast engages the Multiple Demand (MD) network (Fedorenko, Duncan, et al., 2013). A trial consisted of a fixation cross presented for 500 ms, followed by a 3 x 4 grid within which a sequence of blue squares were sequentially flashed at the rate of 1 s per flash (showing one location at a time in the Easy condition, and two at a time in the Hard condition), followed by two sets of locations shown for 1 s, with an additional response time when participants had to decide which set of locations they just saw, and feedback presented for 250 ms (a green checkmark for correct responses or a red cross for incorrect responses), for a total duration of 8 s. A block consisted of 4 trials for a total block duration of 32 s. Each run consisted of 12 experimental blocks (6 per condition) and 4 fixation blocks (16 s each), for a total duration of 448 s (7 min 28 s). The order of conditions varied across runs and participants, but was always palindromic within a run. Each participant performed 2 runs, for a total of 12 blocks per experimental condition.

#### Experiment 7 – Language network localizer

This task was designed to target brain areas that support language processing—i.e., processes related to retrieving words from memory and combining them into structured linguistic representations (Fedorenko et al., 2010; the localizer is available for download from https://evlab.mit.edu/funcloc/). Participants read sentences and lists of disconnected pronounceable nonwords in a blocked design. Although this particular version used written materials, ample evidence has established that near-identical activations are obtained for written and auditory contrasts (e.g., Fedorenko et al., 2010; Scott et al., 2017; Malik-Moraleda, Ayyash et al., 2022; see Fedorenko et al., 2024 for a review). A trial consisted of 12 words/nonwords (presented one at a time, for 450 ms each), preceded by a 100 ms pretrial fixation and followed by a line drawing of a hand pressing a button appearing for 400 ms, and finally a blank screen appearing for 100 ms, for a total trial duration of 6 s. Participants were instructed to press a button when the hand icon appeared, to maintain alertness. A block consisted of 3 trials for a total block duration of 18 s. Each run consisted of 16 experimental blocks (8 per condition) and 5 fixation blocks (14 s each), for a total duration of 358 s (5 min 58 s). The order of conditions varied across runs and participants, but was always palindromic within a run. Each participant performed 2 runs, for a total of 12 blocks per experimental condition.

#### Experiment 8 – Dynamic face perception localizer

This task was designed to target brain areas that specialize for face perception, including dynamic face perception (Kanwisher et al., 1997); Pitcher et al., 2011). Participants watched short video clips of dynamic faces or moving objects (recorded against a black background) in a blocked design. A trial consisted of a single clip and lasted 3 s. A block consisted of 6 trials for a total duration of 18 s. Participants were instructed to watch the movies attentively. Each run consisted of 8 experimental blocks (4 per condition) and 3 rest blocks (18 s each), for a total duration of 198 s (3 min 18 s). During the rest blocks, a series of six uniform color fields were presented for 3 s each (these color fields were designed to maintain the interest of children, for whom this paradigm was originally designed, while approximating a fixation baseline by avoiding any pattern visual input). The order of conditions varied across runs and participants, but was always palindromic within a run. Each participant performed 2 runs, for a total of 8 blocks per experimental condition.

### 4.3 fMRI data acquisition

Whole-brain structural and functional data were collected on a whole-body 3 Tesla Siemens Trio scanner with a 32-channel head coil at the Athinoula A. Martinos Imaging Center at the McGovern Institute for Brain Research at MIT. T1-weighted structural images were collected in 176 axial slices with 1 mm isotropic voxels (repetition time (TR)=2530 ms; echo time (TE)=3.48 ms). Functional, blood oxygenation level-dependent (BOLD) data were acquired using an EPI sequence with a 90° flip angle and using GRAPPA with an acceleration factor of 2; the following parameters were used: thirty-one 4.4 mm thick near-axial slices acquired in an interleaved order (with 10% distance factor), with an in-plane resolution of 2.1 mm × 2.1 mm, FoV in the phase encoding anterior to posterior (A>>P) direction 200 mm and matrix size 96 × 96 voxels, TR=2,000 ms and TE=30 ms. The first 10 s of each run were excluded to allow for steady state magnetization.

### 4.4 fMRI data preprocessing

fMRI data were analyzed using SPM12 (release 7487), CONN EvLab module (release 19b) and other custom MATLAB scripts. Each participant’s functional and structural data were converted from DICOM to NIFTI format. All functional scans were coregistered and resampled using B-spline interpolation to the first scan of the first session (Friston et al., 1995). Potential outlier scans were identified from the resulting subject-motion estimates as well as from BOLD signal indicators using default thresholds in CONN preprocessing pipeline (5 standard deviations above the mean in global BOLD signal change, or framewise displacement values above 0.9 mm, Nieto-Castanon, 2020). Functional and structural data were independently normalized into a common space (the Montreal Neurological Institute [MNI] template; IXI549Space) using SPM12 unified segmentation and normalization procedure (Ashburner & Friston, 2005) with a reference functional image computed as the mean functional data after realignment across all timepoints omitting outlier scans. The output data were resampled to a common bounding box between MNI-space coordinates (-90, -126, -72) and (90, 90, 108), using 2mm isotropic voxels and 4^th^ order spline interpolation for the functional data, and 1mm isotropic voxels and trilinear interpolation for the structural data. Last, the functional data were then smoothed spatially using spatial convolution with a 4 mm FWHM Gaussian kernel.

### 4.4 fMRI data first-level modeling

Effects were estimated using a General Linear Model (GLM) in which each experimental condition was modeled with a boxcar function convolved with the canonical hemodynamic response function (HRF) (fixation was modeled implicitly). Temporal autocorrelations in the BOLD signal timeseries were accounted for by a combination of high-pass filtering with a 128 seconds cutoff, and whitening using an AR(0.2) model (first-order autoregressive model linearized around the coefficient a=0.2) to approximate the observed covariance of the functional data in the context of Restricted Maximum Likelihood estimation (ReML). In addition to main condition effects, other model parameters in the GLM design included first-order temporal derivatives for each condition, modeling spatial variability in the HRF delays, as well as nuisance regressors controlling for the effect of slow linear drifts, subject-motion parameters, and potential outlier scans on the BOLD signal.

### 4.5 Definition of functional regions of interest (fROIs)

To define the fROIs for all the localizer tasks used (prosody, pitch, speech, multiple demand, language, and dynamic faces), we used a group-constrained, subject-specific (GSS) approach (Fedorenko et al., 2010), where a set of binary masks (corresponding to areas of common activation and shown in the main figures for each contrast) is intersected with each participant’s activation map for the relevant contrast. The masks, or parcels, for the multiple demand, language, and dynamic face contrasts come from prior work. For the prosody, pitch, and speech contrasts, the derivation of the masks is described in the next section. Within each mask, a participant-specific fROI was defined as the top 10% of voxels with the highest *t*-values for the localizer contrast (see Lipkin et al., 2022 for evidence that the language fROIs are similar when defined with a fixed statistical threshold). Effect sizes for the critical tasks were estimated in the fROIs by averaging across the voxels within each participant-specific fROI. To estimate the responses to the localizer onditions, a split-half approach was used, so that in each participant half of the data were used to define the fROIs, and the other half—to estimate the effects.

### 4.6 A whole-brain search for areas sensitive to prosody, pitch, and speech

For prosody, as briefly described in the Results, we used a conjunction contrast defined by the sum of the *SP+>SP-* and *NP+>NP-* conditions; for pitch, we used the *Harmonic>Noise* contrast; and for speech, we used the *Nonword>Textures* contrast. Individual activation maps for each contrast were binarized in the following way: voxels that pass the significance threshold of *p*<0.01 for the relevant contrast were denoted as 1 and the rest of the voxels as 0 (note that the use of a relatively liberal threshold at this stage is acceptable because it is only used to create the probabilistic overlap maps, and the statistical tests are performed at a later stage using an independent poartion of the data, as explained below). Individual binarized activation maps were overlaid to create a probabilistic activation overlap map, which was then parcellated using a watershed algorithm, as described in Fedorenko et al. (2010; a custom Matlab toolbox for doing this is available at https://evlab.mit.edu/funcloc) to identify areas of consistent activation across participants. This approach is similar in spirit to the traditional random-effects group analysis, except that it is designed to allow for the use of individual-level fROIs (cf. assuming voxel-wise functional correspondence across participants). The output of the watershed algorithm is a set of masks that correspond to areas of high overlap in the probabilistic map. Parcels that are sufficiently large and show responses in a sufficiently high fraction of the population (see **Figure S1** for details of this procedure for the prosody parcels) are then evaluated using across-runs cross-validation, to ensure that the critical effects are replicable across runs.

For prosody parcels, this approach left us with 18 parcels shown in **Figure S2** (5 left temporal, 4 right temporal parcels 2 left medial-frontal, 2 right medial-frontal, 2 left lateral-frontal and 3 right lateral-frontal, **Figure S2**).

### 4.7 Spatial correlations within and between tasks

To evaluate the similarity of the fine grain voxel activation patterns between the prosody contrast and each of the other contrasts, spatial voxel-wise correlations were computed. Because the activations for all our contrasts are concentrated in particular anatomical areas, for each comparisons, we used a conjunction of the parcels from the relevant contrasts, often maintaining the distinctions between hemispheres and among lobes. For each set of voxels, we correlated the vectors of response (effect size in % BOLD signal change) to the two contrasts using a Fisher-transformed Pearson correlation. For spatial correlations within a task, which evaluated the tasks stability across runs, responses to the task contrast across even and odd runs were computed per participant and then averaged across participants. For the spatial correlations between two tasks, e.g., prosody and task X, 4 spatial correlation values were computed across participant; correlation between odd runs of the prosody task and even runs of task X, and the same for all combinations of odd and even runs across the two tasks. These four values were averaged per participant, and then across participants. For statistical comparison, we compared the spatial correlation value between prosody and task X to the geometric mean of the values within each of the two individual tasks: 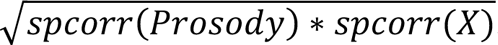.

### 4.8 Probability maps

To create the probability maps (displayed in **Figure 1E** and **Figure 7A**), each participant’s t-map for the relevant localizer contrast was binarized using a top 10% threshold, and the resulting maps were combined such that the value at each voxel reflected the proportion of participants for whom that voxel was among the top 10% most responsive to the localizer contrast.

## Acknowledgements

We would like to thank the Athinoula A. Martinos Imaging Center at the McGovern Institute for Brain Research at MIT, and specifically Steve Shannon and Atsushi Takahashi, as well as all the participants in this study. We also thank Ray Jackendoff, Emalie McMahon and Aaron Wright for their comments on the manuscript, and members of the Fedorenko lab, the audiences at the Neurobiology of Language conference (2022, Philadelphia; 2025, Washington DC), the Cognitive Neuroscience Society meeting (2024, Toronto), as well as Nancy Kanwisher for helpful discussions and comments. TIR was supported by the Poitras Center for Psychiatric Disorders Research at the McGovern Institute for Brain Research, MIT. HK was supported by the Lisa K Yang ICoN graduate fellowship. CC was supported by a graduate fellowship from the Kempner Institute for the Study of Natural and Artificial Intelligence at Harvard University. JSC was supported by NSF grant BCS-1944773. EF was supported by NIH award R01-DC016607, as well as by research funds from MIT’s McGovern Institute for Brain Research, the Siegel Family Quest for Intelligence, and the Department of Brain and Cognitive Sciences Department. This research was supported by a grant from the Simons Foundation to the Simons Center for the Social Brain at MIT.

## Author Contributions

Conceptualization: TIR and EF; Methodology: TIR, HSK, NJ, SS, HK, CC, EF; Software, Validation: TIR, HSK, NJ, SS; Formal analysis: TIR, HSK, NJ; Investigation, Data Curation: TIR, HSK, NJ, SS, HK, CC, EF; Writing-original draft: TIR, EF; Writing-review and editing: TIR, JSC, EF, with additional input from all authors; Visualization: TIR; Supervision: EF; Project administration: EF.

## Supplementary Information

**Figure S1.**
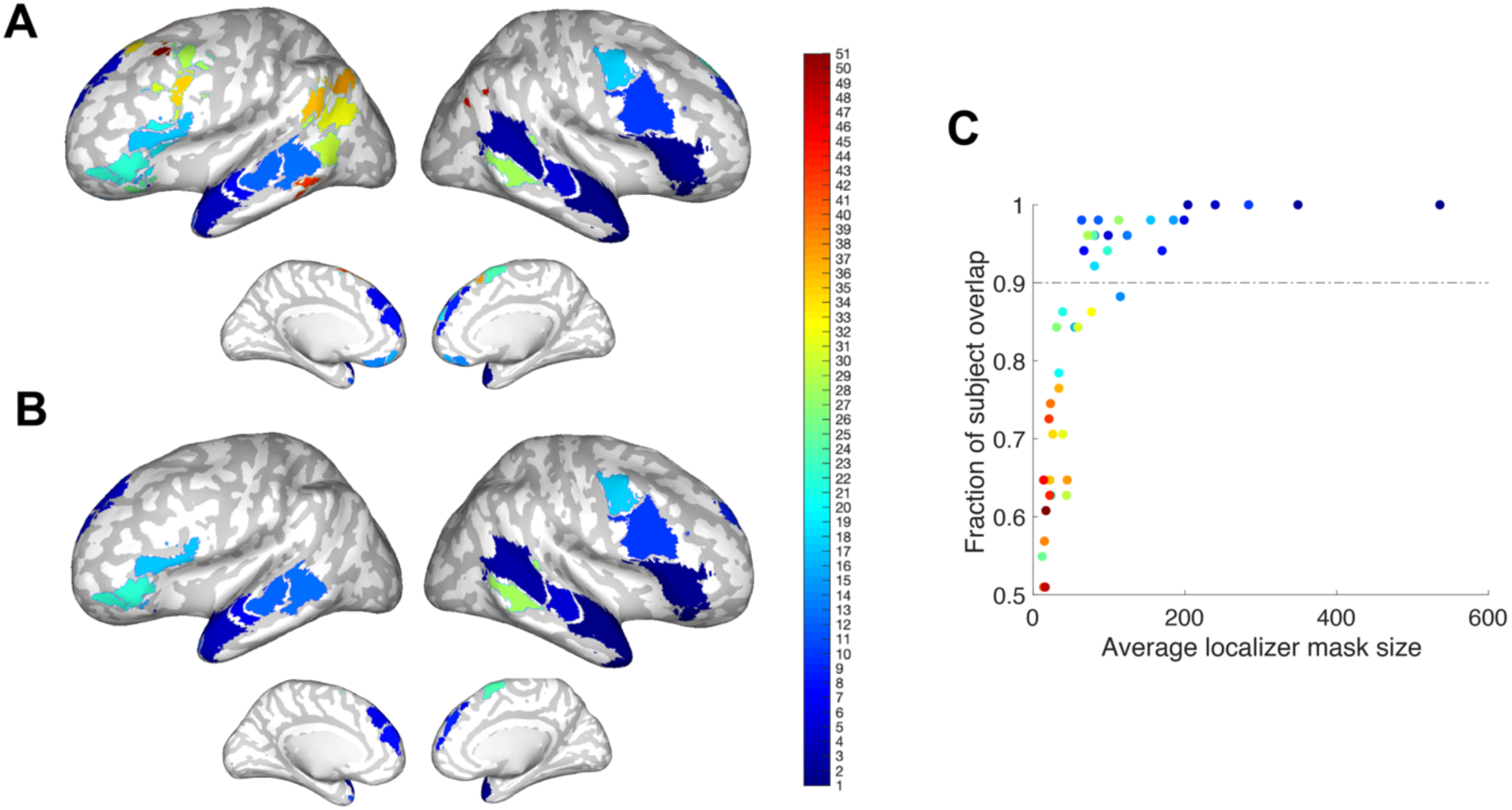
Selection of prosody parcels. **A)** The full set of 51 significant prosody parcels found using our watershed algorithm (<u>Methods</u>). **B**) The reduced set of 18 prosody parcels which were further used in the study. The selection of this reduced set was based on a 90% fraction of participant overlap. **C**) Demonstration of the fraction of participant overlap and the average localizer mask size per each of the full set of parcels (each dot represents a parcel). Parcels above the threshold of 90% (dashed horizontal line) we selected for further analysis.

**Figure S2.**
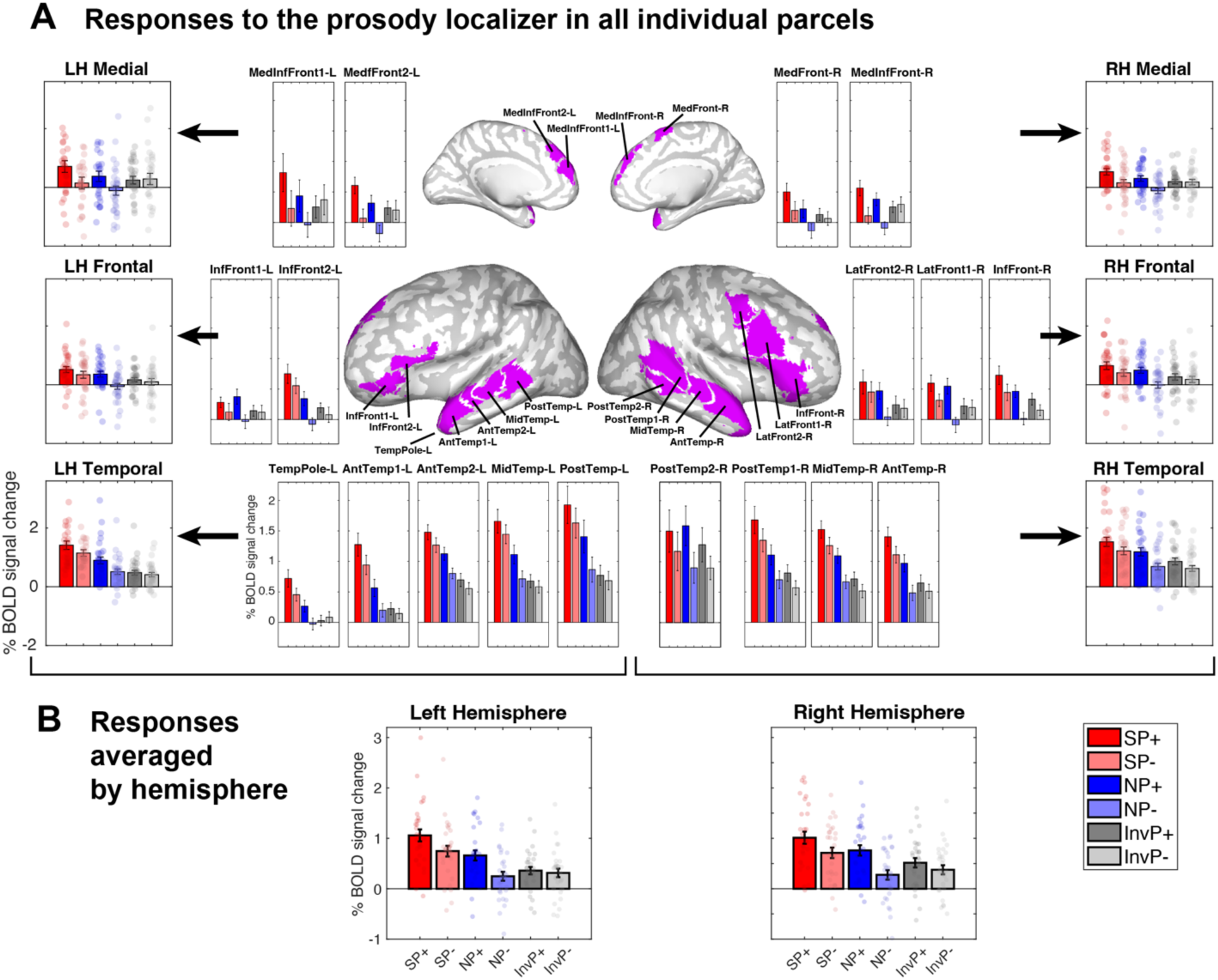
Responses to the prosody localizer in prosody parcels. This figure is similar to Figure 1 in the paper, but the functional response profiles are presented for each of the prosody parcels separately (**A**) and averages per hemisphere are presented (**B**). All technical details are similar to Figure 1.

**Figure S3.**
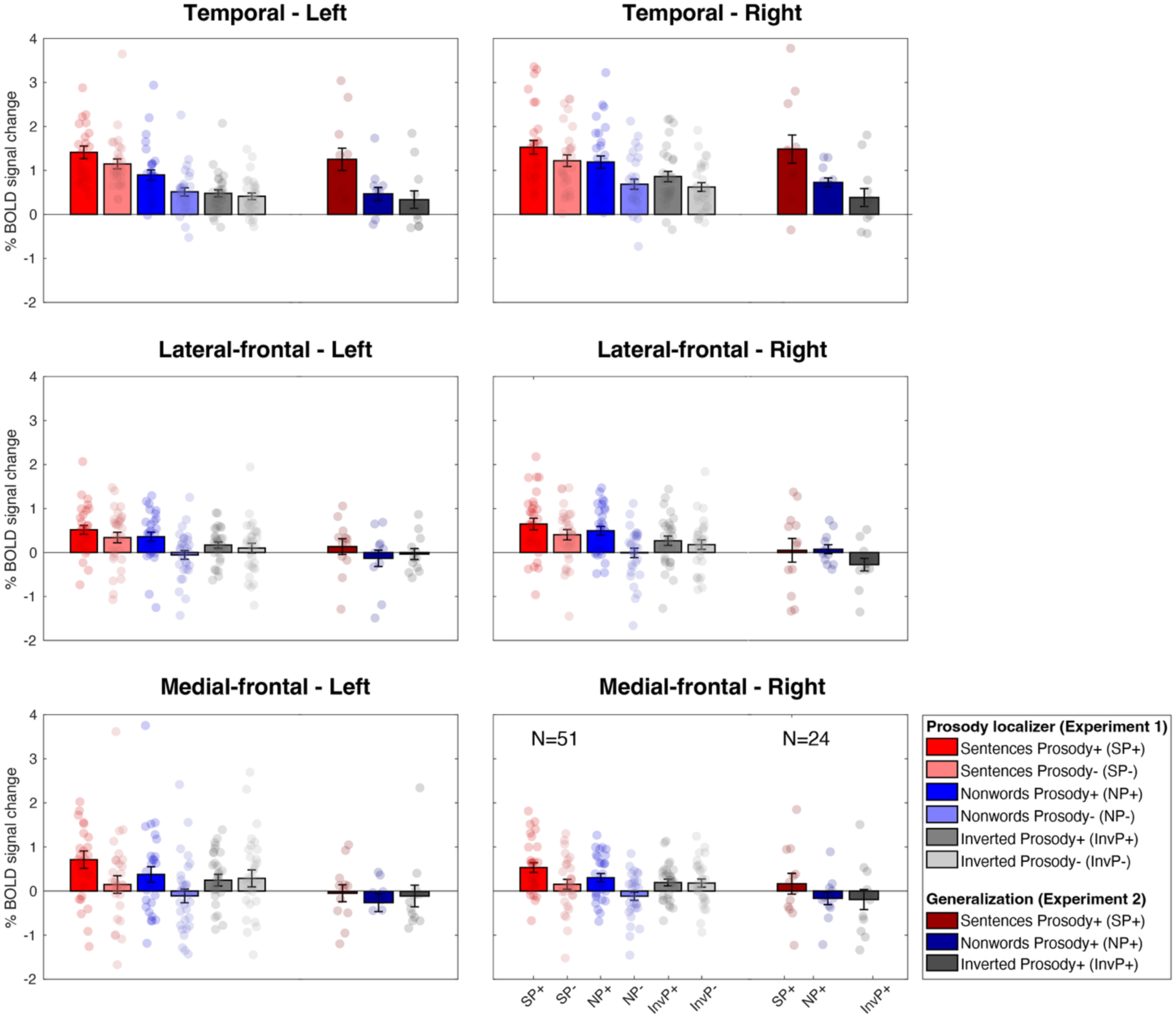
Experiment 2 - Generalization of responses to prosody using naturalistic stimuli. Each subplot represents the average of the specified set of prosody parcels. Within each subplot, the left 6 bars represent the 6 conditions from the prosody localizer (Experiment 1), and the right 3 bars represent the 3 conditions from the generalization experiment (Experiment 2, Methods). Each bar represents the mean response across all individual areas, and across all participants (N=51 in Experiment 1 N=24 in Experiment 2). The bar graphs represent the average (across participants) responses, in % BOLD signal change relative to the fixation baseline. To each condition extracted from individual-level fROIs (error bars are standard error of the mean across participants, and dots correspond to individual participants). Localization and effect size estimation were performed using independent subsets of the data to avoid circularity (Methods).

**Figure S4.**
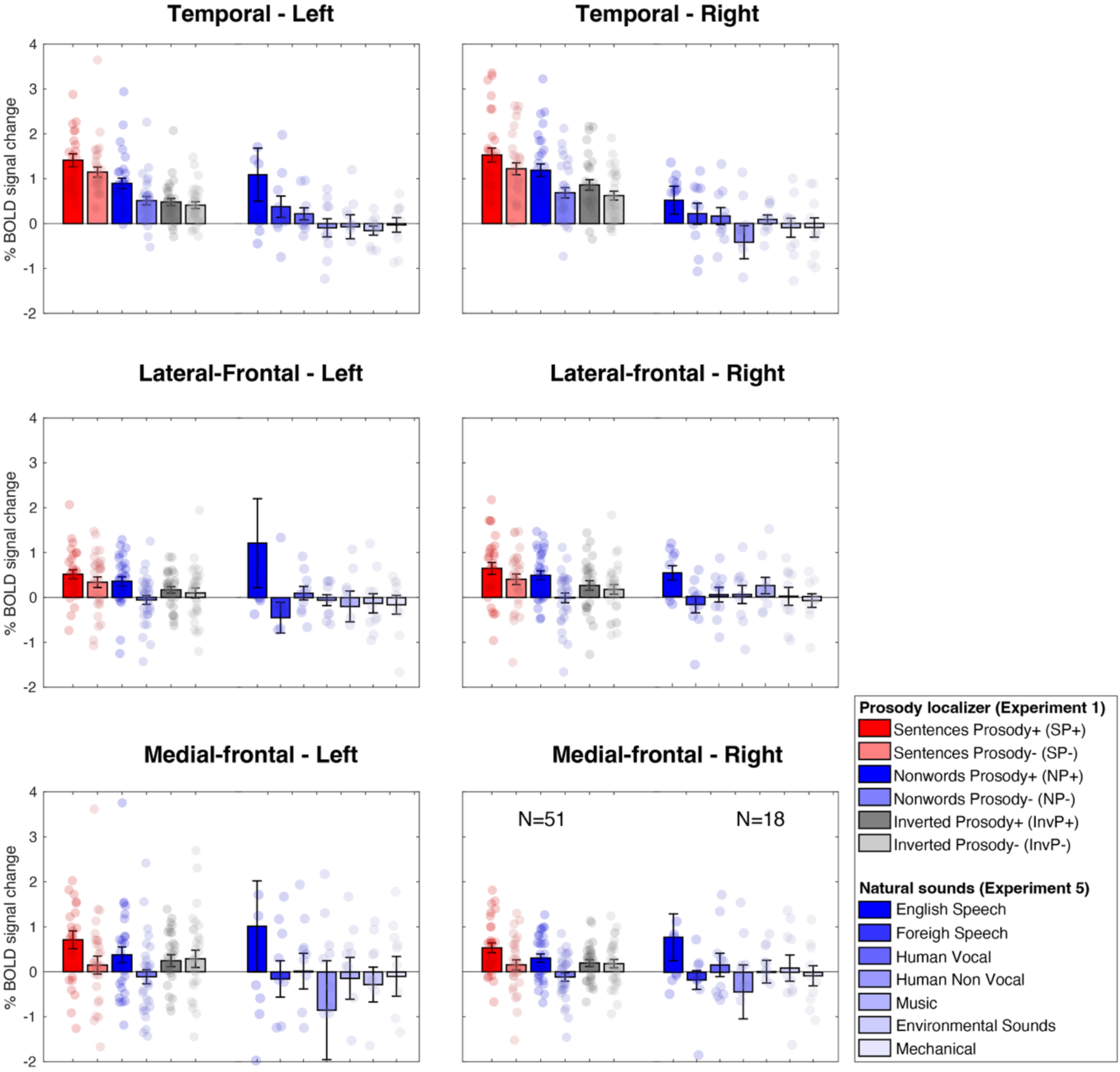
Responses to natural sounds (Experiment 5) in the prosody regions. Each subplot represents the average of the specified set of prosody parcels. Within each subplot, the left 6 bars represent the 6 conditions from the prosody localizer (Experiment 1), and the right 7 bars represent the 7 categories of natural sounds (Experiment 5, Methods). Each bar represents the mean response across all individual areas, and across all participants (N=51 in Experiment 1 N=18 in Experiment 5). The bar graphs represent the average (across participants) responses, in % BOLD signal change relative to the fixation baseline. To each condition extracted from individual-level fROIs (error bars are standard error of the mean across participants, and dots correspond to individual participants). Localization and effect size estimation were performed using independent subsets of the data to avoid circularity (Methods).

**Table S1.** LME results for prosody effects in prosody areas: Distinct LME models are separated by bold titles that specify the effect contrasted in the model and which brain area the model was run for. Each LME model regressed the effect size (% BOLD signal change relative to low level fixation baseline) on a fixed effect variable (Effect), coded as categorical. The first level of the categorical Effect variable was coded as the intercept and the other level(s) as a slope variable, such that the first level of the Effects variable is compared to the fMRI fixation baseline, and any other level of the Effect variable is compared to its first level – the intercept. Thus, the intercept term was of no interest and only the Effect variable was of interest to represent the desired effect. Additional to these fixed effects, the model included random terms where the intercept and effect were grouped by Participant and by ROI (both coded as categorical variables). For example, for the first model, **\*\*Prosody areas LH temporal SP+ vs. SP-**,** the Wilkinson formula was: EffectSize ∼ 1 + Effect + (1+ Effect|Participant) + (1+ Effect|ROI) p<0.001 p<0.01 p<0.05

| Name | Estimate | SE | tStat | DF | pValue | Lower | Upper |
| --- | --- | --- | --- | --- | --- | --- | --- |
| **\*\*Prosody areas LH temporal SP+ vs. SP-\*\*** |  |  |  |  |  |  |  |
| (Intercept – SP+) | 1.42 | 0.199 | 7.15 | 508 | 3.1e-12 | 1.03 | 1.81 |
| Effect – SP- | -0.27 | 0.069 | -3.93 | 508 | 9.6e-05 | -0.41 | -0.14 |
| **\*\*Prosody areas RH Temporal SP+ vs. SP-\*\*** |  |  |  |  |  |  |  |
| (Intercept – SP+) | 1.51 | 0.111 | 13.63 | 406 | 4.4e-35 | 1.29 | 1.72 |
| Effect – SP- | -0.32 | 0.085 | -3.74 | 406 | 2.1e-04 | -0.49 | -0.15 |
| **\*\*Prosody areas LH lateral-frontal SP+ vs. SP-\*\*** |  |  |  |  |  |  |  |
| (Intercept – SP+) | 0.45 | 0.163 | 2.75 | 202 | 0.0064 | 0.13 | 0.77 |
| Effect – SP- | -0.17 | 0.070 | -2.48 | 202 | 0.014 | -0.31 | -0.04 |
| **\*\*Prosody areas RH lateral-frontal SP+ vs. SP-\*\*** |  |  |  |  |  |  |  |
| (Intercept – SP+) | 0.67 | 0.086 | 7.87 | 304 | 6.4e-14 | 0.50 | 0.84 |
| Effect – SP- | -0.24 | 0.055 | -4.26 | 304 | 2.7e-05 | -0.34 | -0.13 |
| **\*\*Prosody areas LH medial-frontal SP+ vs. SP-\*\*** |  |  |  |  |  |  |  |
| (Intercept – SP+) | 0.52 | 0.130 | 4.00 | 202 | 8.8e-05 | 0.26 | 0.77 |
| Effect – SP- | -0.50 | 0.110 | -4.52 | 202 | 1.0e-05 | -0.71 | -0.28 |
| **\*\*Prosody areas RH medial-frontal SP+ vs. SP-\*\*** |  |  |  |  |  |  |  |
| (Intercept – SP+) | 0.59 | 0.085 | 6.92 | 202 | 5.7e-11 | 0.42 | 0.76 |
| Effect – SP- | -0.43 | 0.093 | -4.62 | 202 | 6.8e-06 | -0.61 | -0.25 |
| **\*\*Prosody all areas SP+ vs. SP-\*\*** |  |  |  |  |  |  |  |
| (Intercept – SP+) | 1.01 | 0.137 | 7.38 | 1834 | 2.3e-13 | 0.74 | 1.28 |
| Effect – SP- | -0.31 | 0.047 | -6.49 | 1834 | 1.1e-10 | -0.40 | -0.21 |
| **\*\*Prosody areas LH temporal NP+ vs. NP-\*\*** |  |  |  |  |  |  |  |
| (Intercept – NP+) | 0.85 | 0.190 | 4.46 | 508 | 1.0e-05 | 0.48 | 1.22 |
| Effect – NP- | -0.32 | 0.065 | -4.95 | 508 | 1.0e-06 | -0.45 | -0.19 |
| **\*\*Prosody areas RH Temporal NP+ vs. NP-\*\*** |  |  |  |  |  |  |  |
| (Intercept – NP+) | 1.05 | 0.109 | 9.59 | 406 | 9.2e-20 | 0.83 | 1.27 |
| Effect – NP- | -0.39 | 0.077 | -5.06 | 406 | 6.4e-07 | -0.54 | -0.24 |
| **\*\*Prosody areas LH lateral-frontal NP+ vs. NP-\*\*** |  |  |  |  |  |  |  |
| (Intercept – NP+) | 0.30 | 0.072 | 4.10 | 202 | 5.9e-05 | 0.15 | 0.44 |
| Effect – NP- | -0.32 | 0.086 | -3.68 | 202 | 3.0e-04 | -0.48 | -0.15 |
| **\*\*Prosody areas RH lateral-frontal NP+ vs. NP-\*\*** |  |  |  |  |  |  |  |
| (Intercept – NP+) | 0.43 | 0.075 | 5.74 | 304 | 2.3e-08 | 0.28 | 0.58 |
| Effect – NP- | -0.39 | 0.073 | -5.41 | 304 | 1.3e-07 | -0.54 | -0.25 |
| **\*\*Prosody areas LH medial-frontal NP+ vs. NP-\*\*** |  |  |  |  |  |  |  |
| (Intercept – NP+) | 0.17 | 0.115 | 1.44 | 202 | 0.15 | -0.06 | 0.39 |
| Effect – NP- | -0.25 | 0.109 | -2.28 | 202 | 0.024 | -0.46 | -0.03 |
| **\*\*Prosody areas RH medial-frontal NP+ vs. NP-\*\*** |  |  |  |  |  |  |  |
| (Intercept – NP+) | 0.24 | 0.069 | 3.39 | 202 | 8.3e-04 | 0.10 | 0.37 |
| Effect – NP- | -0.29 | 0.075 | -3.86 | 202 | 1.5e-04 | -0.44 | -0.14 |
| **\*\*Prosody all areas NP+ vs. NP-\*\*** |  |  |  |  |  |  |  |
| (Intercept – NP+) | 0.62 | 0.113 | 5.45 | 1834 | 5.8e-08 | 0.40 | 0.84 |
| Effect – NP- | -0.34 | 0.062 | -5.42 | 1834 | 6.7e-08 | -0.46 | -0.21 |
| **\*\*Prosody areas LH temporal InvP+ vs. InvP-\*\*** |  |  |  |  |  |  |  |
| (Intercept – InvP+) | 0.46 | 0.134 | 3.45 | 508 | 6.2e-04 | 0.20 | 0.73 |
| Effect – InvP- | -0.07 | 0.048 | -1.43 | 508 | 0.15 | -0.16 | 0.03 |
| **\*\*Prosody areas RH Temporal InvP+ vs. InvP-\*\*** |  |  |  |  |  |  |  |
| (Intercept – InvP+) | 0.75 | 0.092 | 8.07 | 406 | 8.1e-15 | 0.56 | 0.93 |
| Effect – InvP- | -0.22 | 0.066 | -3.35 | 406 | 8.8e-04 | -0.35 | -0.09 |
| **\*\*Prosody areas LH lateral-frontal InvP+ vs. InvP-\*\*** |  |  |  |  |  |  |  |
| (Intercept – InvP+) | 0.11 | 0.051 | 2.11 | 202 | 0.036 | 0.01 | 0.21 |
| Effect – InvP- | -0.05 | 0.064 | -0.82 | 202 | 0.41 | -0.18 | 0.07 |
| **\*\*Prosody areas RH lateral-frontal InvP+ vs. InvP-\*\*** |  |  |  |  |  |  |  |
| (Intercept – InvP+) | 0.22 | 0.066 | 3.33 | 304 | 9.9e-04 | 0.09 | 0.35 |
| Effect – InvP- | -0.14 | 0.065 | -2.11 | 304 | 0.036 | -0.27 | -0.01 |
| **\*\*Prosody areas LH medial-frontal InvP+ vs. InvP-\*\*** |  |  |  |  |  |  |  |
| (Intercept – InvP+) | 0.13 | 0.086 | 1.53 | 202 | 0.13 | -0.04 | 0.30 |
| Effect – InvP- | 0.05 | 0.114 | 0.45 | 202 | 0.65 | -0.17 | 0.28 |
| **\*\*Prosody areas RH medial-frontal InvP+ vs. InvP-\*\*** |  |  |  |  |  |  |  |
| (Intercept – InvP+) | 0.17 | 0.057 | 3.06 | 202 | 0.0025 | 0.06 | 0.29 |
| Effect – InvP- | -0.08 | 0.071 | -1.11 | 202 | 0.27 | -0.22 | 0.06 |
| **\*\*Prosody all areas InvP+ vs. InvP-\*\*** |  |  |  |  |  |  |  |
| (Intercept – InvP+) | 0.38 | 0.082 | 4.57 | 1834 | 5.1e-06 | 0.22 | 0.54 |
| Effect – InvP- | -0.10 | 0.051 | -1.96 | 1834 | 0.0503 | -0.20 | 0.00 |

**Table S2.** LME results for intelligibility effects in prosody areas: Details are similar to Table S1 p<0.001 p<0.01 p<0.05

| Name | Estimate | SE | tStat | DF | pValue | Lower | Upper |
| --- | --- | --- | --- | --- | --- | --- | --- |
| **\*\*Prosody areas LH temporal SP+ vs. NP+\*\*** |  |  |  |  |  |  |  |
| (Intercept – SP+) | 1.42 | 0.197 | 7.20 | 508 | 2.2e-12 | 1.03 | 1.81 |
| Effect – NP+ | -0.57 | 0.070 | -8.14 | 508 | 3.0e-15 | -0.71 | -0.43 |
| **\*\*Prosody areas RH Temporal SP+ vs. NP+\*\*** |  |  |  |  |  |  |  |
| (Intercept – SP+) | 1.51 | 0.108 | 13.99 | 406 | 1.4e-36 | 1.29 | 1.72 |
| Effect – NP+ | -0.46 | 0.091 | -5.00 | 406 | 8.6e-07 | -0.64 | -0.28 |
| **\*\*Prosody areas LH lateral-frontal SP+ vs. NP+\*\*** |  |  |  |  |  |  |  |
| (Intercept – SP+) | 0.45 | 0.157 | 2.86 | 202 | 0.0047 | 0.14 | 0.76 |
| Effect – NP+ | -0.15 | 0.159 | -0.97 | 202 | 0.33 | -0.47 | 0.16 |
| **\*\*Prosody areas RH lateral-frontal SP+ vs. NP+\*\*** |  |  |  |  |  |  |  |
| (Intercept – SP+) | 0.67 | 0.085 | 7.89 | 304 | 5.4e-14 | 0.51 | 0.84 |
| Effect – NP+ | -0.24 | 0.058 | -4.19 | 304 | 3.7e-05 | -0.36 | -0.13 |
| **\*\*Prosody areas LH medial-frontal SP+ vs. NP+\*\*** |  |  |  |  |  |  |  |
| (Intercept – SP+) | 0.52 | 0.131 | 3.96 | 202 | 1.0e-04 | 0.26 | 0.78 |
| Effect – NP+ | -0.35 | 0.104 | -3.41 | 202 | 7.9e-04 | -0.56 | -0.15 |
| **\*\*Prosody areas RH medial-frontal SP+ vs. NP+\*\*** |  |  |  |  |  |  |  |
| (Intercept – SP+) | 0.59 | 0.084 | 7.06 | 202 | 2.6e-11 | 0.42 | 0.75 |
| Effect – NP+ | -0.35 | 0.083 | -4.27 | 202 | 3.0e-05 | -0.52 | -0.19 |
| **\*\*Prosody areas LH temporal SP- vs. NP-\*\*** |  |  |  |  |  |  |  |
| (Intercept – SP-) | 1.15 | 0.191 | 6.02 | 508 | 3.4e-09 | 0.77 | 1.52 |
| Effect – NP- | -0.62 | 0.068 | -9.04 | 508 | 3.2e-18 | -0.75 | -0.48 |
| **\*\*Prosody areas RH Temporal SP- vs. NP-\*\*** |  |  |  |  |  |  |  |
| (Intercept – SP-) | 1.19 | 0.091 | 13.02 | 406 | 1.3e-32 | 1.01 | 1.37 |
| Effect – NP- | -0.53 | 0.076 | -6.92 | 406 | 1.8e-11 | -0.68 | -0.38 |
| **\*\*Prosody areas LH lateral-frontal SP- vs. NP-\*\*** |  |  |  |  |  |  |  |
| (Intercept – SP-) | 0.28 | 0.153 | 1.81 | 202 | 0.072 | -0.03 | 0.58 |
| Effect – NP- | -0.30 | 0.159 | -1.86 | 202 | 0.065 | -0.61 | 0.02 |
| **\*\*Prosody areas RH lateral-frontal SP- vs. NP-\*\*** |  |  |  |  |  |  |  |
| (Intercept – SP-) | 0.44 | 0.075 | 5.83 | 304 | 1.4e-08 | 0.29 | 0.58 |
| Effect – NP- | -0.40 | 0.069 | -5.86 | 304 | 1.2e-08 | -0.54 | -0.27 |
| **\*\*Prosody areas LH medial-frontal SP- vs. NP-\*\*** |  |  |  |  |  |  |  |
| (Intercept – SP-) | 0.02 | 0.128 | 0.18 | 202 | 0.86 | -0.23 | 0.28 |
| Effect – NP- | -0.10 | 0.115 | -0.91 | 202 | 0.37 | -0.33 | 0.12 |
| **\*\*Prosody areas RH medial-frontal SP- vs. NP-\*\*** |  |  |  |  |  |  |  |
| (Intercept – SP-) | 0.16 | 0.102 | 1.58 | 202 | 0.12 | -0.04 | 0.36 |
| Effect – NP- | -0.21 | 0.100 | -2.14 | 202 | 0.034 | -0.41 | -0.02 |

**Table S3.**
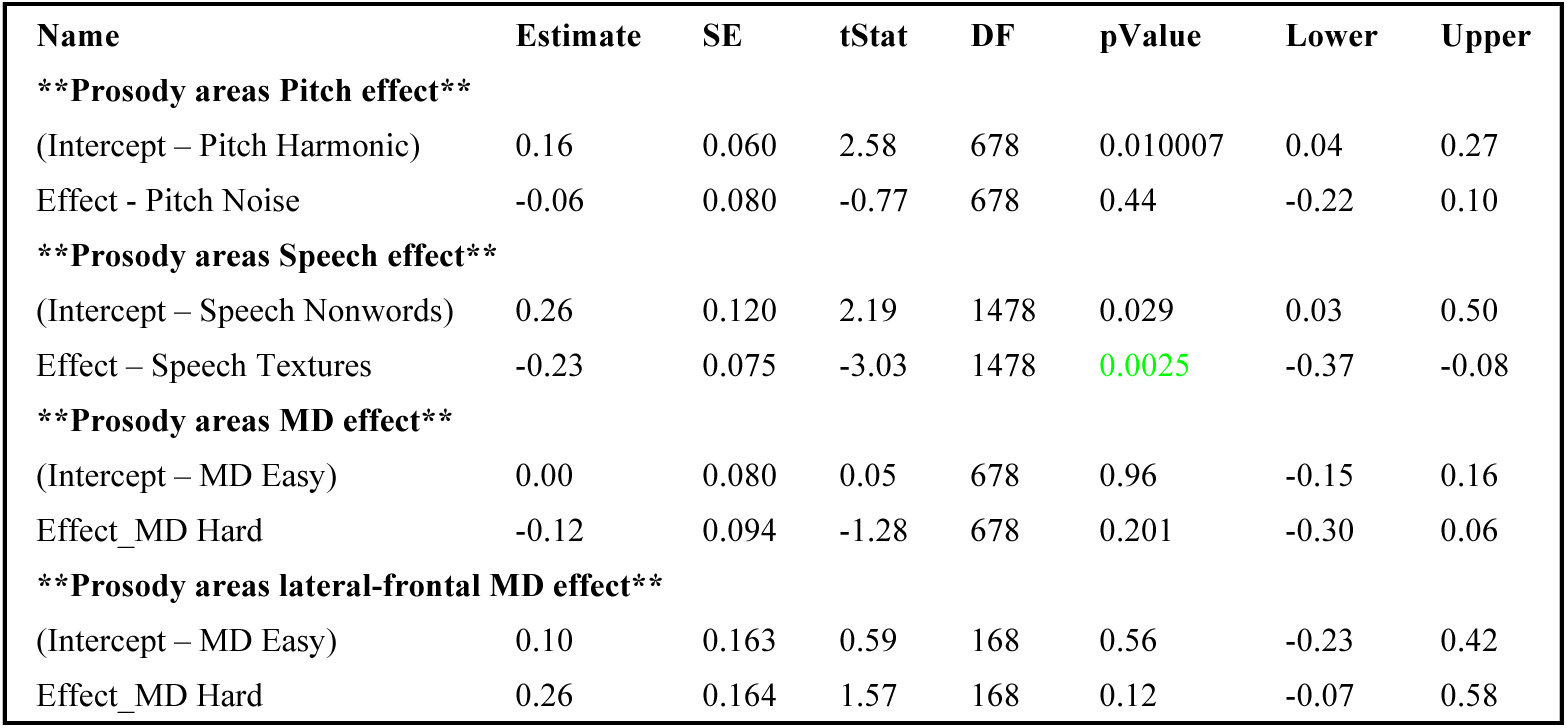
LME results for other functional effects in prosody areas: Details are similar to Table S1 p<0.001 p<0.01 p<0.05

| Name | Estimate | SE | tStat | DF | pValue | Lower | Upper |
| --- | --- | --- | --- | --- | --- | --- | --- |
| **\*\*Prosody areas Pitch effect\*\*** |  |  |  |  |  |  |  |
| (Intercept – Pitch Harmonic) | 0.16 | 0.060 | 2.58 | 678 | 0.010007 | 0.04 | 0.27 |
| Effect - Pitch Noise | -0.06 | 0.080 | -0.77 | 678 | 0.44 | -0.22 | 0.10 |
| **\*\*Prosody areas Speech effect\*\*** |  |  |  |  |  |  |  |
| (Intercept – Speech Nonwords) | 0.26 | 0.120 | 2.19 | 1478 | 0.029 | 0.03 | 0.50 |
| Effect – Speech Textures | -0.23 | 0.075 | -3.03 | 1478 | 0.0025 | -0.37 | -0.08 |
| **\*\*Prosody areas MD effect\*\*** |  |  |  |  |  |  |  |
| (Intercept – MD Easy) | 0.00 | 0.080 | 0.05 | 678 | 0.96 | -0.15 | 0.16 |
| Effect_MD Hard | -0.12 | 0.094 | -1.28 | 678 | 0.201 | -0.30 | 0.06 |
| **\*\*Prosody areas lateral-frontal MD effect\*\*** |  |  |  |  |  |  |  |
| (Intercept – MD Easy) | 0.10 | 0.163 | 0.59 | 168 | 0.56 | -0.23 | 0.42 |
| Effect_MD Hard | 0.26 | 0.164 | 1.57 | 168 | 0.12 | -0.07 | 0.58 |

**Table S4.** LME results for prosody effects in other functional areas: Details are similar to Table S1 p<0.001 p<0.01 p<0.05

| Name | Estimate | SE | tStat | DF | pValue | Lower | Upper |
| --- | --- | --- | --- | --- | --- | --- | --- |
| **Pitch areas prosody effect (SP+,NP+ vs. SP-, NP-)** |  |  |  |  |  |  |  |
| (Intercept – Pros+ [SP+,NP+]) | 3.96 | 0.380 | 10.41 | 134 | 5.7e-19 | 3.21 | 4.71 |
| Effect – Pros- [SP-, NP-] | -0.09 | 0.133 | -0.68 | 134 | 0.5 | -0.35 | 0.17 |
| **Pitch areas prosody conditions relative to SP+** |  |  |  |  |  |  |  |
| (Intercept – SP+) | 3.63 | 0.349 | 10.40 | 198 | 1.7e-20 | 2.94 | 4.32 |
| Effect – SP- | 0.26 | 0.174 | 1.49 | 198 | 0.14 | -0.08 | 0.60 |
| Effect – NP+ | 0.66 | 0.184 | 3.59 | 198 | 4.2e-04 | 0.30 | 1.03 |
| Effect – NP- | 0.22 | 0.181 | 1.23 | 198 | 0.22 | -0.14 | 0.58 |
| Effect – InP+ | 0.37 | 0.200 | 1.85 | 198 | 0.065 | -0.02 | 0.77 |
| Effect – InvP- | 0.16 | 0.175 | 0.92 | 198 | 0.36 | -0.18 | 0.51 |
| **Pitch areas prosody conditions relative to NP+** |  |  |  |  |  |  |  |
| (Intercept – NP+) | 4.29 | 0.413 | 10.38 | 198 | 2.0e-20 | 3.48 | 5.11 |
| Effect – NP- | -0.44 | 0.183 | -2.40 | 198 | 0.017 | -0.80 | -0.08 |
| Effect – SP+ | -0.66 | 0.184 | -3.59 | 198 | 4.2e-04 | -1.03 | -0.30 |
| Effect – SP- | -0.40 | 0.176 | -2.29 | 198 | 0.023 | -0.75 | -0.06 |
| Effect – InvP+ | -0.29 | 0.210 | -1.39 | 198 | 0.17 | -0.70 | 0.12 |
| Effect – InvP- | -0.50 | 0.176 | -2.84 | 198 | 0.0049 | -0.85 | -0.15 |
| **Speech areas prosody effect (SP+,NP+ vs. SP-, NP-)** |  |  |  |  |  |  |  |
| (Intercept – Pros+ [SP+, NP+]) | 3.85 | 0.148 | 25.94 | 286 | 4.3e-77 | 3.56 | 4.14 |
| Effect – Pros- [SP-, NP-] | -0.10 | 0.083 | -1.19 | 286 | 0.24 | -0.26 | 0.06 |
| **Speech areas prosody conditions relative to SP+** |  |  |  |  |  |  |  |
| (Intercept – SP+) | 3.78 | 0.156 | 24.27 | 426 | 2.5e-82 | 3.48 | 4.09 |
| Effect – SP- | 0.20 | 0.114 | 1.76 | 426 | 0.08 | -0.02 | 0.42 |
| Effect – NP+ | 0.13 | 0.105 | 1.26 | 426 | 0.209 | -0.07 | 0.34 |
| Effect – NP- | -0.27 | 0.109 | -2.44 | 426 | 0.015 | -0.48 | -0.05 |
| Effect – InvP+ | -0.80 | 0.177 | -4.52 | 426 | 8.0e-06 | -1.15 | -0.45 |
| Effect – InvP- | -1.04 | 0.130 | -7.95 | 426 | 1.8e-14 | -1.29 | -0.78 |
| **Speech areas prosody conditions relative to NP+** |  |  |  |  |  |  |  |
| (Intercept – NP+) | 3.91 | 0.156 | 25.14 | 426 | 3.4e-86 | 3.61 | 4.22 |
| Effect – NP- | -0.40 | 0.105 | -3.78 | 426 | 1.8e-04 | -0.60 | -0.19 |
| Effect – SP+ | -0.13 | 0.105 | -1.26 | 426 | 0.209 | -0.34 | 0.07 |
| Effect – SP- | 0.07 | 0.107 | 0.64 | 426 | 0.52 | -0.14 | 0.28 |
| Effect – InvP+ | -0.93 | 0.192 | -4.85 | 426 | 1.8e-06 | -1.31 | -0.55 |
| Effect – InvP- | -1.17 | 0.142 | -8.24 | 426 | 2.1e-15 | -1.45 | -0.89 |
| **Speech areas prosody conditions relative to NP-** |  |  |  |  |  |  |  |
| (Intercept – NP-) | 3.52 | 0.141 | 24.96 | 426 | 2.3e-85 | 3.24 | 3.79 |
| Effect – NP+ | 0.40 | 0.105 | 3.78 | 426 | 1.8e-04 | 0.19 | 0.60 |
| Effect – SP+ | 0.27 | 0.109 | 2.44 | 426 | 0.015 | 0.05 | 0.48 |
| Effect – SP- | 0.47 | 0.109 | 4.29 | 426 | 2.2e-05 | 0.25 | 0.68 |
| Effect – InvP+ | -0.53 | 0.197 | -2.71 | 426 | 0.00704 | -0.92 | -0.15 |
| Effect – InvP- | -0.77 | 0.145 | -5.33 | 426 | 1.6e-07 | -1.06 | -0.49 |
| **Speech areas prosody inversion effect** |  |  |  |  |  |  |  |
| (Intercept – Inverted speech [InvP+, InvP-]) | 2.86 | 0.155 | 18.53 | 430 | 8.7e-57 | 2.56 | 3.17 |
| Effect – Forward speech [SP+, SP-, NP+, NP-] | 0.93 | 0.146 | 6.41 | 430 | 3.9e-10 | 0.65 | 1.22 |
| **Language LH areas SP+ vs. SP-** |  |  |  |  |  |  |  |
| (Intercept – SP+) | 2.09 | 0.244 | 8.57 | 508 | 1.2e-16 | 1.61 | 2.57 |
| Effect – SP- | -0.11 | 0.078 | -1.35 | 508 | 0.18 | -0.26 | 0.05 |
| **Language RH areas SP+ vs. SP-** |  |  |  |  |  |  |  |
| (Intercept – SP+) | 1.59 | 0.249 | 6.37 | 508 | 4.1e-10 | 1.10 | 2.07 |
| Effect – SP- | -0.25 | 0.064 | -3.88 | 508 | 1.2e-04 | -0.38 | -0.12 |
| **\*\*Language LH areas NP+ vs. NP-\*\*** |  |  |  |  |  |  |  |
| (Intercept – NP+) | 1.05 | 0.210 | 4.98 | 508 | 8.9e-07 | 0.63 | 1.46 |
| Effect – NP- | -0.42 | 0.079 | -5.37 | 508 | 1.2e-07 | -0.58 | -0.27 |
| **\*\*Language RH areas NP+ vs. NP-\*\*** |  |  |  |  |  |  |  |
| (Intercept – NP+) | 1.00 | 0.200 | 4.99 | 508 | 8.3e-07 | 0.61 | 1.39 |
| Effect – NP- | -0.35 | 0.060 | -5.79 | 508 | 1.2e-08 | -0.47 | -0.23 |

**Table S5.**
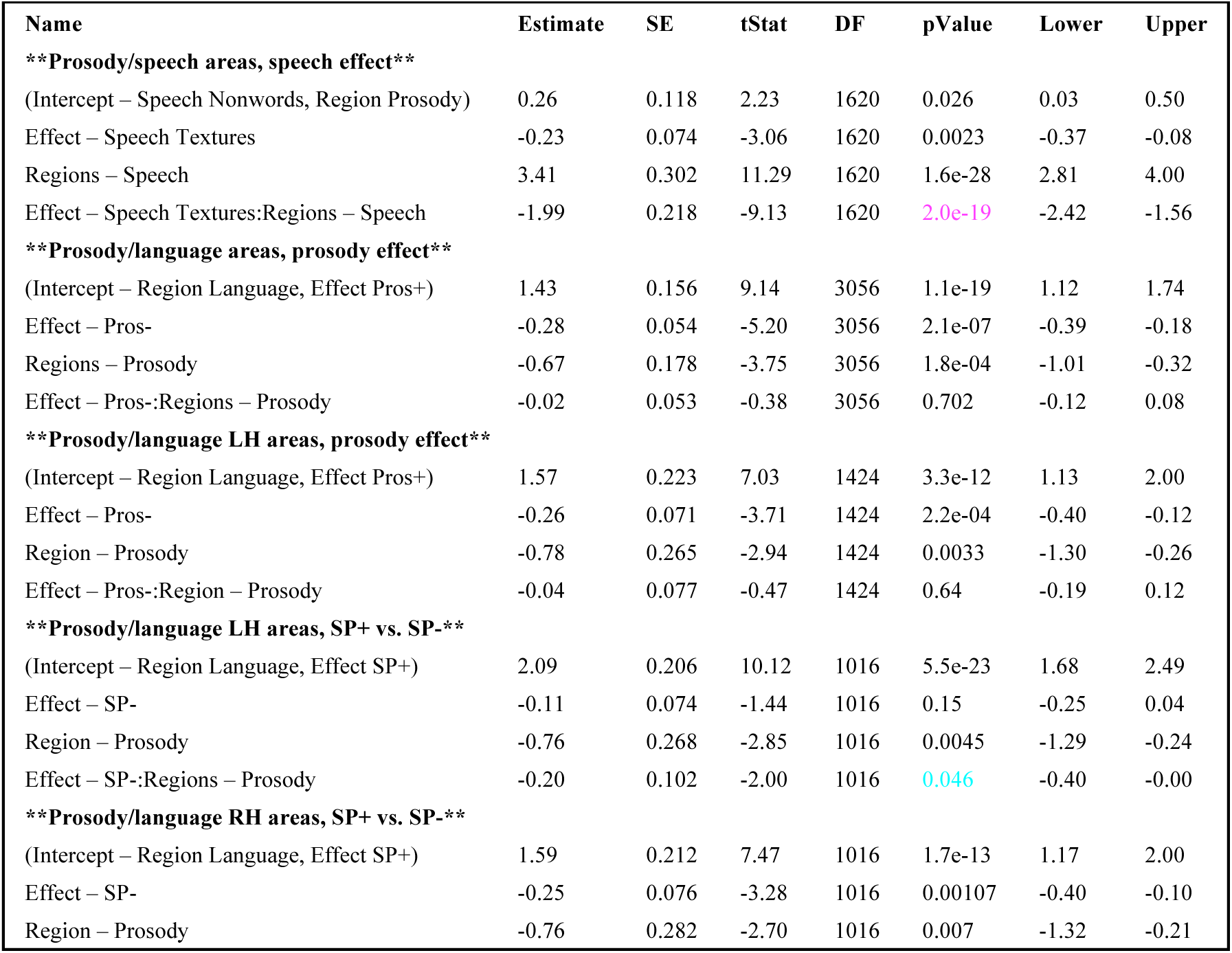
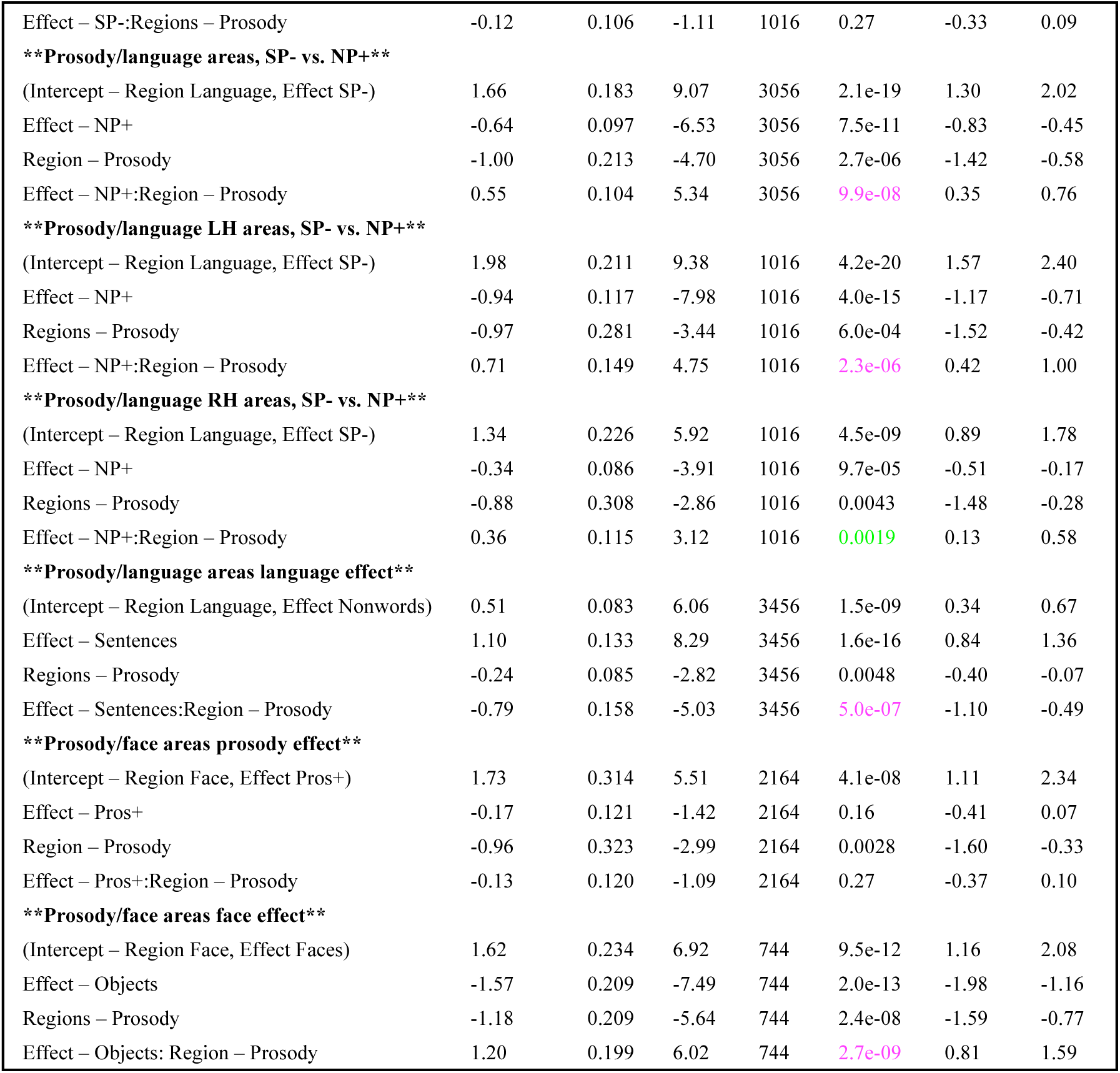
LME results for interactions between area type (prosody or other areas) with functional effects: Details are similar to Table S1, except that: Each LME model regressed the effect size on three fixed effects variables; Effect, Region (two options for brain regions, e.g. prosody and language), and their interaction. Both variables were coded as categorical and thus, the intercept represented the combination of the first levels of Effect and of Region. The other level(s) of Effect and Region were coded as slope variable, which were compared to the intercept. Finally, the multiplication of both the slope variables modeled the interaction of the two variables. Thus, the intercept as well as the main effects of each variable are of no interest and only the interaction term was of interest here. Additional to these fixed effects, the model included random terms where the intercept and effect (but not region) were grouped by Participant and by ROI (both coded as categorical variables). For example, for the first model, **\*\*Prosody/speech areas, speech effect**,** the Wilkinson formula was: EffectSize ∼ 1 + Effect*Region + (1+ Effect|Participant) + (1+ Effect|ROI) which is equivalent to: EffectSize ∼ 1 + Effect + Region + Effect:Region + (1+ Effect|Participant) + (1+ Effect|ROI). p<0.001 p<0.01 p<0.05

**Table S6.**
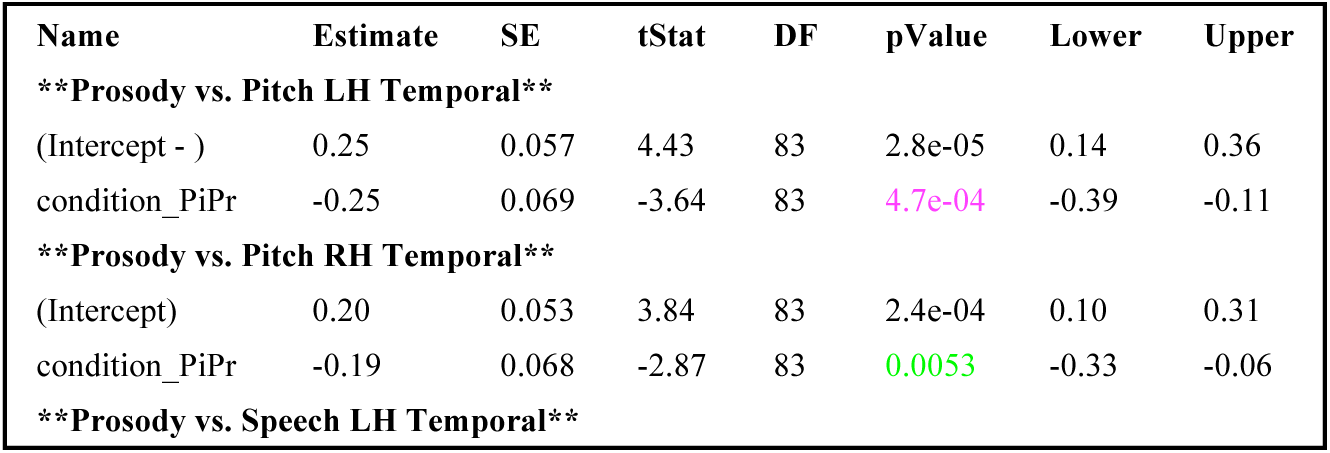
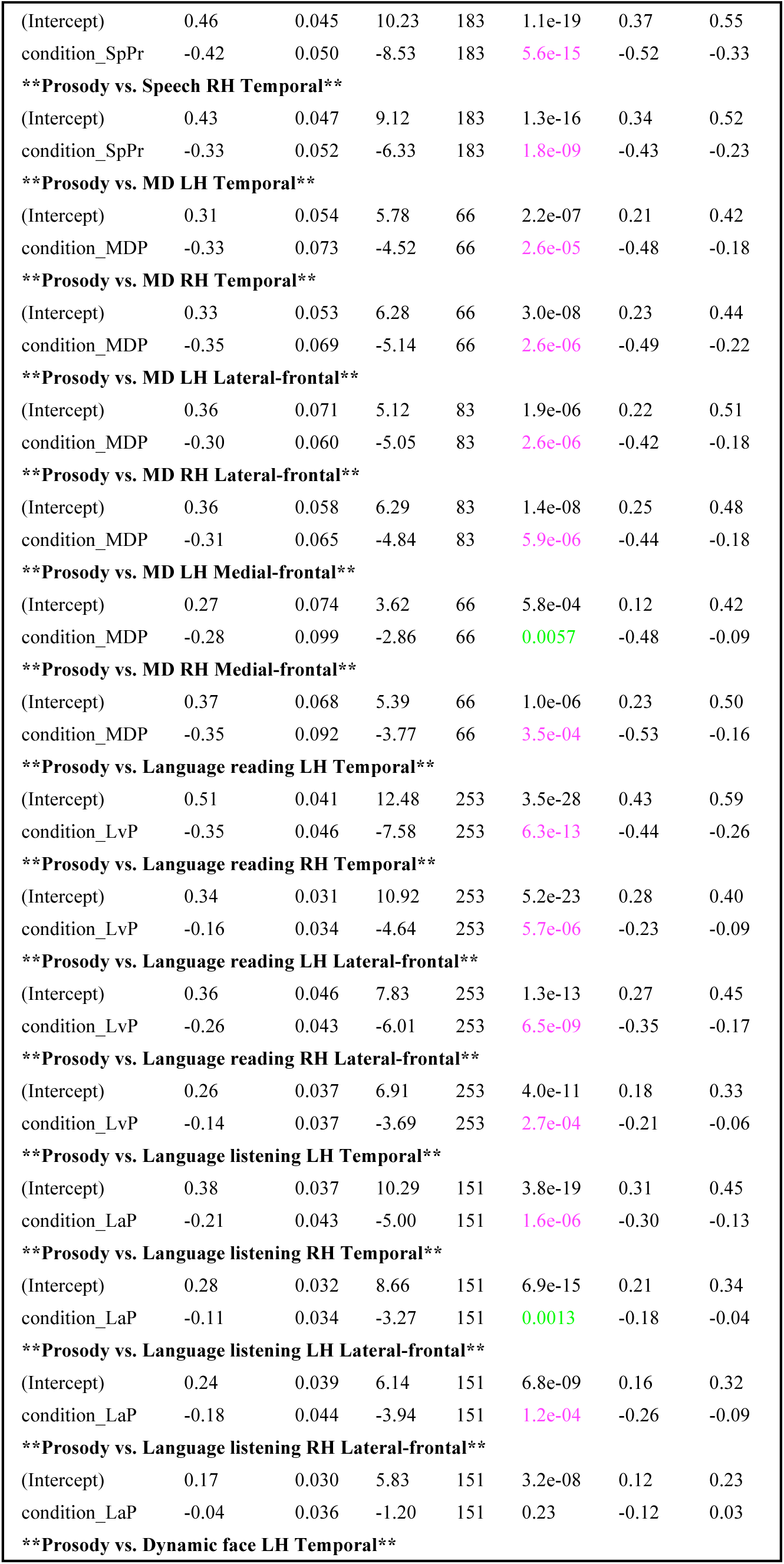
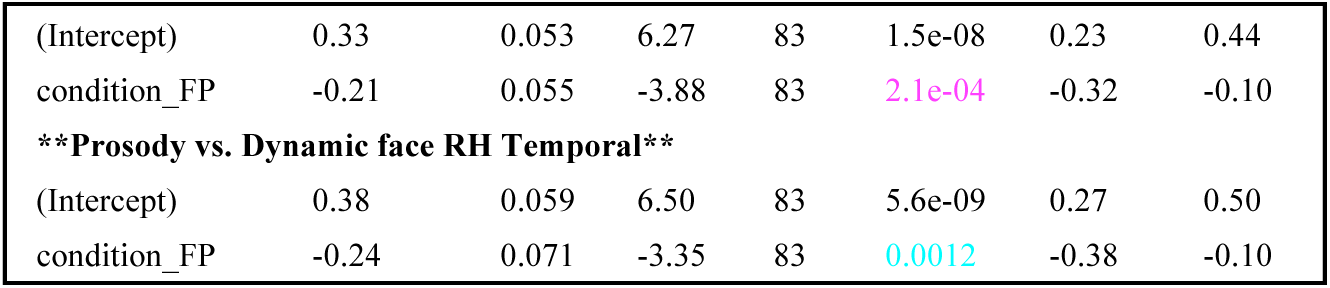
LME results for spatial correlations between prosody and all other functional areas: P value map: p<0.001 p<0.01 p<0.05

